# PAD2 knockout reduces myelin protein aggregates, modulates neuroinflammation and protects motor neurons, axons and neuromuscular junction in a SOD1-ALS mouse model

**DOI:** 10.64898/2026.05.30.729013

**Authors:** Issa O. Yusuf, Rogerio L.A. Silva, George G. Amoako, Paul R. Thompson, Zuoshang Xu

**Affiliations:** Department of Biochemistry and Molecular Biotechnology, University of Massachusetts Chan Medical School, Worcester, MA, USA; Program in Chemical Biology, University of Massachusetts Chan Medical School, Worcester, MA 01605 USA

**Author notes:** Corresponding authors: Issa O. Yusuf, Department of Biochemistry and Molecular Biotechnology, University of Massachusetts Chan Medical School, 364 Plantation St, Worcester, MA 01605, USA, Zuoshang Xu, Department of Biochemistry and Molecular Biotechnology, University of Massachusetts Chan Medical School, 364 Plantation St, Worcester, MA 01605, USA.

**Keywords:** PAD2 knockout, protein citrullination, neurodegeneration, deimination, neurodegenerative diseases, myelin degeneration, protein aggregation

## Abstract

**Background:** Dysregulated peptidyl deiminase 2 (PAD2) and aberrant protein citrullination (PC), a posttranslational modification (PTM), are involved in various inflammatory and neurodegenerative diseases. We previously showed in transgenic mice and postmortem human tissues that PC and PAD2 are altered in amyotrophic lateral sclerosis (ALS), a neurodegenerative disease characterized by motor neurons loss, paralysis, and death. Herein, we investigated the role of PAD2 in ALS by PAD2 knockout in a SOD1-ALS mouse model.

**Methods:** To investigate the role of PAD2-induced citrullination in ALS pathogenesis, we generated PAD2 knockout (PAD2KO) in SOD1^G93A^ ALS mouse model and investigated the consequent modulation on the neuropathology and clinical symptoms, using molecular biology techniques such as qPCR, Western blotting, confocal microscopy, and electron microscopy. Additionally, we identified C3 as being citrullinated in human ALS using ionFinder.

**Results:** Our results show that PAD2KO blocked the increased PC and reduced myelin basic protein (MBP) aggregates in the ALS model. PAD2KO also improved motor neuron survival and the integrity of myelin, axons, and neuromuscular junctions, and reduced microgliosis in the white matter and C3 protein levels in astrocytes. Clinically, data from monitoring the body weight changes suggests that PAD2KO modulates the course of the disease in the ALS mouse model, accelerating the onset while slowing the progression after the onset, and modestly extending the survival of male mice.

**Conclusion:** These results show that PAD2 is responsible for the increased PC in ALS and PC contributes to neuroinflammation and degeneration of motor neurons and myelinated axons. The modest modulation of the disease phenotype suggests that the role of PC in ALS is complex, involving altered PC in numerous proteins and in multiple cell types. Future studies are needed to investigate how PC modulates individual protein functions in various cell types to understand the contribution of PC to ALS pathogenesis.

## Background

Amyotrophic lateral sclerosis (ALS) is a progressive neurodegenerative disorder characterized by motor neuron degeneration, muscle wasting, paralysis, and death. Approximately 90 percent of ALS cases are sporadic, whereas ∼10 percent are familial (1). Although the precise pathological mechanisms remain elusive, several processes, including genetic mutations, protein aggregation, abnormal RNA processing, cytoskeletal disruption, mitochondrial dysfunction, endoplasmic reticulum (ER) stress, reactive oxygen species, neuroinflammation, non-cell autonomous toxicity, and white matter degeneration, are reportedly involved in ALS disease pathogenesis (2–5). Despite considerable advances, the disease still defies a cure.

Protein citrullination (PC) or deimination, is an irreversible posttranslational modification (PTM) that converts peptidyl-arginine to peptidyl-citrulline (6). PC is catalyzed by a family of enzymes known as protein arginine deiminases (PADs). PAD activity is tightly regulated by calcium levels (7). At high calcium levels (1–10 mM), PADs are active and catalyze PC. This reaction leads to a loss of positive charges, thereby impacting ionic and hydrogen bond formation. PADs are differentially expressed in tissues, and they have shared as well as distinct substrate specificities. In the CNS, PAD2 is the most dominantly expressed PAD. Aberrant PAD2 activity and PC are implicated in diverse pathologies, including neurodegenerative conditions such as Alzheimer’s disease (AD), Parkinson’s disease (PD), prion disease, multiple sclerosis (MS), ischemic and traumatic brain injuries (8–13). Targeting PAD2 has been reported to ameliorate the neurodegeneration phenotypes in some disease models (14–19).

We recently investigated the patterns of PAD2 expression and PC changes in ALS mouse models. These studies unveiled that PAD2 and PC change dynamically during ALS disease progression (20). After disease onset, PAD2 expression increases, culminating in the accumulation of PC in disease-impacted areas such as the spinal cord. Investigation at the cellular level revealed a nuanced pattern. PAD2 and PC increase in astrocytes but decrease in neurons during disease progression. A later study in human ALS verified these observations (21). In this study, we investigated the role of PAD2 in ALS by crossing PAD2 knockout (PAD2KO) mice with the SOD1^G93A^ ALS mouse model. We show that PAD2KO blocked the increase in PC observed during disease progression and reduced citrullinated protein and MBP aggregates. At the cellular level, PAD2KO improved motor neuron survival, down-regulated complement C3 levels in astrocytes, reduced microgliosis in the white matter, protected myelin and axonal integrity, and slowed denervation at neuromuscular junction (NMJ). Clinically, PAD2KO modulated the course of the disease, modestly accelerating disease onset while slowing disease progression after onset. PAD2KO also increased the lifespan in the SOD1^G93A^ mouse model, albeit only in males. These results demonstrate that PAD2 is responsible for the increase in PC in ALS and suggest that PAD2 and PC contribute to ALS pathogenesis in a complex manner depending on the ensemble effects from various citrullination events among many proteins.

## Materials and methods

### Transgenic mice

The high expression line of transgenic mice expressing human mutant SOD1G93A (B6SJL-Tg(SOD1*G93A)1Gur/J) Stock No: 002726, were purchased from Jackson Lab (Bar Harbor, ME), and bred onto FVB background for more than ten generations. Non-transgenic (nTg) littermates of SOD^G93A^ were used as controls. PAD2 knockout (PAD2KO) mice on FVB/NJ background were generously provided by Scott Coonrod (Cornell University, Ithaca, New York, USA). We crossed SOD^G93A^ mice with PAD2KO mice to generate SOD1^G93A^/PAD2KO mice. Mice were maintained at the University of Massachusetts Medical School animal facility according to the guidelines provided by the Institutional Animal Care and Use Committee (IACUC).

### Behavioral analysis

Mouse body weight was measured weekly, and their overall health was monitored daily towards the paralysis stage of the disease. The mice reach paralysis or end stage when one of the following conditions is met: (1) two or more limbs become paralyzed (completely immobile), and (2) incapable of righting itself in 10s when placed on either side. The paralysis stage was the endpoint of the experiment, and the mouse was sacrificed for tissue harvesting.

### RNA isolation and quantitative real-time PCR

Total RNA was extracted from lumber spinal cord tissue using (Trizol™ Reagent; ThermoFisher Scientific, 15596018), and cDNA synthesis performed using iScript cDNA Synthesis Kit (Bio-Rad, 1708891) according to the manufacturer’s protocol. Quantitative PCR (qPCR) was performed using SsoFast™ EvaGreen® Supermix (Bio-Rad, 1725200). CFX96™ Touch Real-Time PCR detection system was used with the following program: 2 min of pre-denaturation at 95 °C, 45 consecutive cycles of 5 s denaturation at 95 °C, 20 s annealing at 60 °C. Relative mRNA expression was normalized to 60S ribosomal protein L32 (RPL-32). See supplementary Table 1 for the list of primers used.

### Western blotting

Mice were decapitated under deep anesthesia. Tissues were quickly harvested, snap-frozen in liquid N_2_, and stored at –80 °C. Frozen tissues were weighed, homogenized in homogenization buffer [25 mM phosphate pH 7.6, 5 mM EDTA, 1% (vol/vol) SDS, 0.5% (vol/vol) Triton X-100, 0.5% (vol/vol) deoxycholic acid, protease and phosphatase inhibitor cocktail (Thermo Scientific)], and then centrifuged at 16,060 x g for 10 min at 4 °C. The supernatant was collected as protein sample. Protein concentration was measured using Bradford assay (Bio-Rad, 5000006). The samples were heated in Laemmli buffer (Bio-Rad 1610747) plus 10% β-Mercaptoethanol (Bio-Rad, 1610710) at 95 °C for 10 min, and equal amounts of protein were electrophoretically separated by SDS-PAGE (Bio-Rad) and transferred to PVDF membranes (Thermo Scientific, 88518). The membranes were blocked with 5% (wt/vol) nonfat dry milk (Boston Bioproducts Inc., P-1400) in PBS for 1 h at RT, and incubated with the appropriate primary antibody (supplementary Table 2) overnight at 4 °C. After washing with phosphate-buffered saline with 0.1% Tween 20 (PBST) three times for 5 min each, membranes were incubated with horseradish peroxidase (HRP)-conjugated secondary antibodies (supplementary Table 3), for 1 h at RT. Membranes washed three times for 5 min each and signals were visualized by SuperSignal West Pico PLUS Chemiluminescent Substrate (Thermo Scientific, 34580) reagent and detected by the Amersham Imager 600 (GE).

### Immunofluorescence and immunohistochemistry

Anesthetized mice were transcardially perfused with cold PBS followed by 4% (wt/vol) paraformaldehyde (PFA) in PBS. The perfused mice were then immersed in 4% PFA at 4 °C for another 24–48 h. Spinal cord tissues were dissected and immersed in PBS containing 30% (wt/vol) sucrose at 4 °C for 2–3 days, then frozen in optimal cutting temperature (OCT) freezing media (Sakura) and stored at –80 °C. Frozen sections were cut at 20 μm using a cryostat. Sections were incubated in blocking solution [5% (vol/vol) donkey serum (Sigma, D9663), 0.15% Triton-X100 and (2%) nonfat dry milk in PBS, pH 7.4] for 1 h at RT. For mouse-on-mouse staining, anti-mouse IgG Fab fragments (Jackson Immuuno Research Laboratories, Inc., 115-007-003) were added to the blocking solution. Following the blocking step, sections were incubated with appropriate primary antibody (Supplementary Table 2) in the blocking solution overnight at 4 °C, washed with PBST three times for 10 min each and incubated in the appropriate secondary antibody (Supplementary Table 3) at RT for 2 h. For immunofluorescence, the sections were washed three times and mounted on super frost plus slide (Fisher scientific, 15-188-48) using Vectashield mounting medium containing DAPI (Vector Laboratories, H-1800) and sealed with nail polish. Images of the spinal cord sections were taken with a confocal microscope (Leica). For immunohistochemistry, sections were washed three times in PBS containing 0.25% (vol/vol) Tween 20, stained following the manufacturer’s protocol for Vectastain ABC kit, Elite PK-6100 standard, ImmPact DAB peroxidase Substrate kit (Vector Lab, SK-4105), and counterstained with Hematoxylene (Vector lab, H-3401). The sections were then mounted on slides and dried overnight at 55 °C. After soaking in Xylene twice for 2 min each, the slides were sealed with Permount mounting medium (Fisher Chemical SP15-100). Images were taken with a Nikon Optiphot microscope equipped with a SPOT Insight 2.0 Mp Firewire Color digital camera.

### Detection of protein citrullination by AMC method

Protein citrullination was detected using the anti-modified citrulline (AMC) detection kit (Millipore, 17-347B) according to manufacturer’s protocol. For detection of citrullinated proteins on immunoblots, proteins were resolved by SDS-PAGE and transferred to a PVDF membrane. The membrane was washed twice with tap water and incubated with AMC reagent at 37 °C overnight without agitation. The modified membranes were rinsed 4–5 times with tap water, blocked in freshly prepared 5% nonfat dry milk in TBST for 1 h at RT with constant agitation, incubated with up to 10 mL of a 1:1000 dilution of AMC antibody (Millipore, MABS487) for 2 h at RT, washed three times with TBST for 10 min each, incubated with goat anti-Human IgG HRP-conjugate in TBST-Milk (1:2000) for 1 h at RT, and washed three times with TBST for 10 min each. Finally, the membranes were rinsed one time with water, and the staining signal was visualized as described above (Western blotting). For immunostaining to detect citrullinated proteins, the method of Asaga and Senshu was adapted (22). This procedure is similar to the immunostaining described previously except that tissue section is treated with AMC reagent as described above. The sections were washed three times for 10 min each, blocked for 1 h at RT, incubated with primary antibody (AMC, 1:100) overnight at RT. The sections were then incubated with secondary antibody for 2 h at RT. Sections were mounted on slides and taken for confocal microscopy imaging.

### Sedimentation assays

Mouse lumber spinal cords were homogenized using a handheld polytron for 20 s in lysis buffer [50 mM Tris·HCl, 150 mM NaCl, 0.5% deoxycholic acid, 1% (vol/vol) Triton X-100, 5 mM EDTA] with protease and phosphatase inhibitor cocktail (1:100 dilution, Thermo Scientific™, 78442). Homogenates were centrifuged at 12,000 × g at 4 °C for 5 min. The supernatants were moved to another tube, and protein concentration was determined using Bradford assay (Bio-rad). The pellets were rinsed three times with PBS and dissolved with one-half the volume of the homogenate: 50µl Laemmli buffer by heating to 95 °C for 10 min, briefly sonicated, and then cleared by centrifugation. 80 µg of supernatant protein and three times equivalent volume of the pellet from each sample were resolved by SDS-PAGE, and proteins transferred to a PVDF membrane. Western blots for detecting proteins were as described above.

### Filter trap assays

Preparation and protein concentration measurement of lumber spinal cord homogenate are described in the sedimentation assay section. 200 µg of protein were equalized with lysis buffer to the same volume and then diluted with 20 volumes of PBS (pH 7.4) containing 1% (vol/vol) SDS. The solution was sonicated (70 W, 50% output, 30 s) and then filtered with a vacuum through PVDF membranes (0.45 mm pore size) using a 96-well dot-blot apparatus (Schleicher and Schuell, Inc.). Each well was washed two times with PBS. The membrane was removed from the device and rinsed in water. Detection of citrullinated proteins and other proteins are as described above (Western blotting).

### Visualization and quantification of motor neurons, myelin and axonal degeneration

For quantification of ventral horn motor neurons, lumbar enlargement of the fixed spinal cords was sectioned on a cryostat at 20 μm thickness. The sections were stained with goat anti-ChAT antibody to reveal motor neurons as described in immunohistochemistry procedure. Motor neuron numbers in the ventral horn region were counted manually from each section. For visualization of spinal root axons, mice were fixed by transcardial perfusion using 4% (wt/vol) paraformaldehyde and 2.5% (wt/vol) glutaraldehyde in 0.1 M sodium cacodylate (pH 7.6). Tissues were further fixed by soaking in the same fixative at 4 °C for 48 h. Spinal cord and L5 roots attached to dorsal root ganglia were dissected and postfixed with 1% (wt/vol) osmium tetroxide in 0.1 M sodium cacodylate (pH 7.6) for 1 h at RT. Samples were washed three times with deionized H_2_O for 10 min, dehydrated through a graded series of ethanol (10, 30, 50, 70, 85, 95% for 20 min each) and to three changes of 100% ethanol. Samples were infiltrated first with two changes of 100% Propylene Oxide and then with a 50%/50% propylene oxide / SPI-Pon 812 resin mixture. On the following day, five changes of fresh 100% SPI-Pon 812 resin were performed before the samples were polymerized at 68 °C in embedding molds. The samples were trimmed for TEM. 1 µm semi-thin sections were placed on glass slides and stained with toluidine blue, examined and photographed by light microscopy using a Nikon Optiphot microscope equipped with a SPOT Insight 2.0 Mp Firewire Color digital camera. For quantification of intact and degenerating axons, we manually counted using ImageJ Multi-point tool. 10 to 12 image frames were analyzed from each animal and were averaged. The values from each animal in every group were further averaged and compared statistically. For TEM imaging, 100 nm thin sections were placed on copper support grids and contrasted with Lead citrate and Uranyl acetate. Sections were examined using the CM10 with 100 Kv accelerating voltage and images were captured using a Gatan TEM CCD camera.

### Visualization and quantification of neuromuscular junctions

Mice were anesthetized and transcardially perfused with cold PBS followed by 4% (wt/vol) paraformaldehyde (PFA) in PBS. The perfused mice were then immersed in 4% PFA at 4 °C for another 24–48 h. After fixation, gastrocnemius muscles were dissected out and immersed in PBS containing 30% (wt/vol) sucrose at 4 °C for 2–3 days. Muscles were embedded in OCT medium, frozen rapidly in OCT and stored at –80 °C. Frozen sections were cut at 20 μm using a cryostat, placed on Superfrost Plus slides, and stored at –20 °C. Slides were allowed to defrost for 30 min before use. Slides were washed three times with PBS for 5 min and then sections were blocked using blocking solution [5% (vol/vol) normal serum in PBS, pH 7.4] for 1 h at RT and then incubated with a primary antibody in the blocking solution overnight at 4 °C. A primary antibody solution of Rabbit anti-synaptophysin (ThermoFisher, 1:200) and Chicken anti-NFL (Novus Biologicals, 1:200) were applied. Slides were washed three times with PBS for 5 min each. A secondary antibody solution of Alexa-568nm-labelled Donkey anti-Rabbit (ThermoFisher, 1:500), Alexa-647nm-labelled Donkey anti-Chicken (ThermoFisher, 1:200), and Alexa-488nm α-Bungarotoxin (ThermoFisher, 1:200) diluted in blocking solution were applied overnight at 4°C in the dark. Slides were imaged on a confocal microscope (Leica). Neuromuscular junctions (NMJ) were then quantified by two methods: NMJ occupancy and NMJ contact. NMJ occupancy was calculated using the formula: NMJ Occupancy (%) = (Presynaptic Area/Postsynaptic Area) * 100. To quantify postsynaptic NMJ contacted by the presynaptic terminal we counted the number of a-bungarotoxin-stained postsynaptic terminals that have an overlap with synaptophysin signal and multiplied by 100.

### Quantitative image analysis

The parameters of staining and imaging were identical for all images taken in each study. ImageJ was used to process and assess all images. For quantification of PC, PAD2, GFAP, Iba1, Cd68, MBP, NF-L, and C3, the staining intensity was measured from 6-10 image frames in each animal and were averaged. The number of animals in each group is in figure legends. The values from each animal in each group were further averaged and compared statistically. For quantification of Olig2 and Ng2 positive cell numbers, we manually counted the cells using ImageJ Multi-point tool. The numbers from 20 to 30 image frames in each animal were averaged and the average was used to represent the number of cells in each animal. For quantification of aggregates, confocal images were taken from ventral and ventrolateral spinal cord white matter. Because the aggregates have very high staining intensity, images were taken at relatively low exposure to exclude background and low intensity signals. A cut-off for aggregate size was set at 10 µm^2^. Signals from other cellular processes, such as astrocytes and myelin, were excluded by applying a shape factor filter to include only structures with shape circularity of 0.4–1. The few remaining linear processes were eliminated manually. The numbers from 10 to 12 image frames in each animal were averaged and the average was used to represent the number of aggregates in each animal. The number of aggregates were normalized to the area of 1 mm^2^.

### Data mining for the C3 citrullination site

Raw mass spectrometry data (syn53424735) (23) were processed using FragPipe (version 22.0) with the TMT18 workflow and adding citrullination as a variable modification. Peptide-spectrum match (PSM) output files were analyzed using ionFinder (24) to identify citrullination sites and assign “true” citrullination events. In parallel, the raw PSM-level TMT intensities were log₂-transformed and median-normalized across all TMT plexes. These normalized values were subsequently aggregated to the protein or peptide level and used for downstream quantitative analyses. The data were mined from 7 healthy controls, 7 Sporadic ALS, 1 C9ORF72 asymptomatic and 2 C9ORF72 symptomatic cases. For statistical analysis, control group comprises healthy controls while the ALS group comprises Sporadic ALS and the C9ORF72 cases.

### Statistics

Results are shown as mean ± standard error of mean (SEM). GraphPad Prism (GraphPad Software Inc. vr. 8.4.3) was used to analyze statistical differences. One-way ANOVA with Bonferroni post hoc test was used in comparing the means of more than two groups. Log-rank test was used to compare the Kaplan-Meier survival curves. Statistical significance was set at p<0.05.

## Results

### PAD2 knockout prevents increased protein citrullination (PC) and citrullinated protein aggregates in the SOD1^G93A^ ALS model

To determine the role of PAD2 in ALS, we crossed SOD1^G93A^ mice with PAD2KO mice. We confirmed the lack of PAD2 expression in SOD1^G93A^/PAD2KO mice by qPCR, Western blots, and immunofluorescence staining (Fig. S1). To determine whether PAD2 knockout results in compensatory changes in other PADs expressed in CNS, we quantified PAD1, PAD3 and PAD4 in spinal cords by qPCR, and Western blotting. PAD1 mRNA levels were exceedingly low in mouse spinal cord and could not be reliably measured by qPCR. PAD3 and PAD4 levels were not altered by PAD2KO in either qPCR or Western blotting measurements (Fig. S2). PAD4 has been reportedly upregulated in neurons in AD and PD, and in myelin in MS (25). However, our attempt in measuring the level of PAD4 in specific cell types was thwarted by a lack of PAD4-specific antibodies (Fig. S3).

To determine how PAD2 impacts PC in ALS, we performed Western blots for citrullinated proteins. As reported previously (20), PC increased dramatically in SOD1^G93A^ mice. PAD2KO prevented this increase (Fig. 1A, B). Immunofluorescent (IF) staining for citrullinated proteins in the spinal cord confirmed this finding (Fig. 1C-E). Importantly, the citrullinated protein aggregates that were abundant in the white matter of SOD1^G93A^ mice became undetectable in SOD1^G93A^/PAD2KO mice (Fig. 1C, E). To confirm the absence of the aggregates, we performed filter trap assay using spinal cord lysate. Aggregated citrullinated proteins were abundant in SOD1^G93A^ mice but undetectable in SOD1^G93A^/PAD2KO mice (Fig. 1F, G). Thus, PAD2 activity was responsible for the increase of PC and citrullinated protein aggregates in this ALS mouse model.

**Figure 1.**
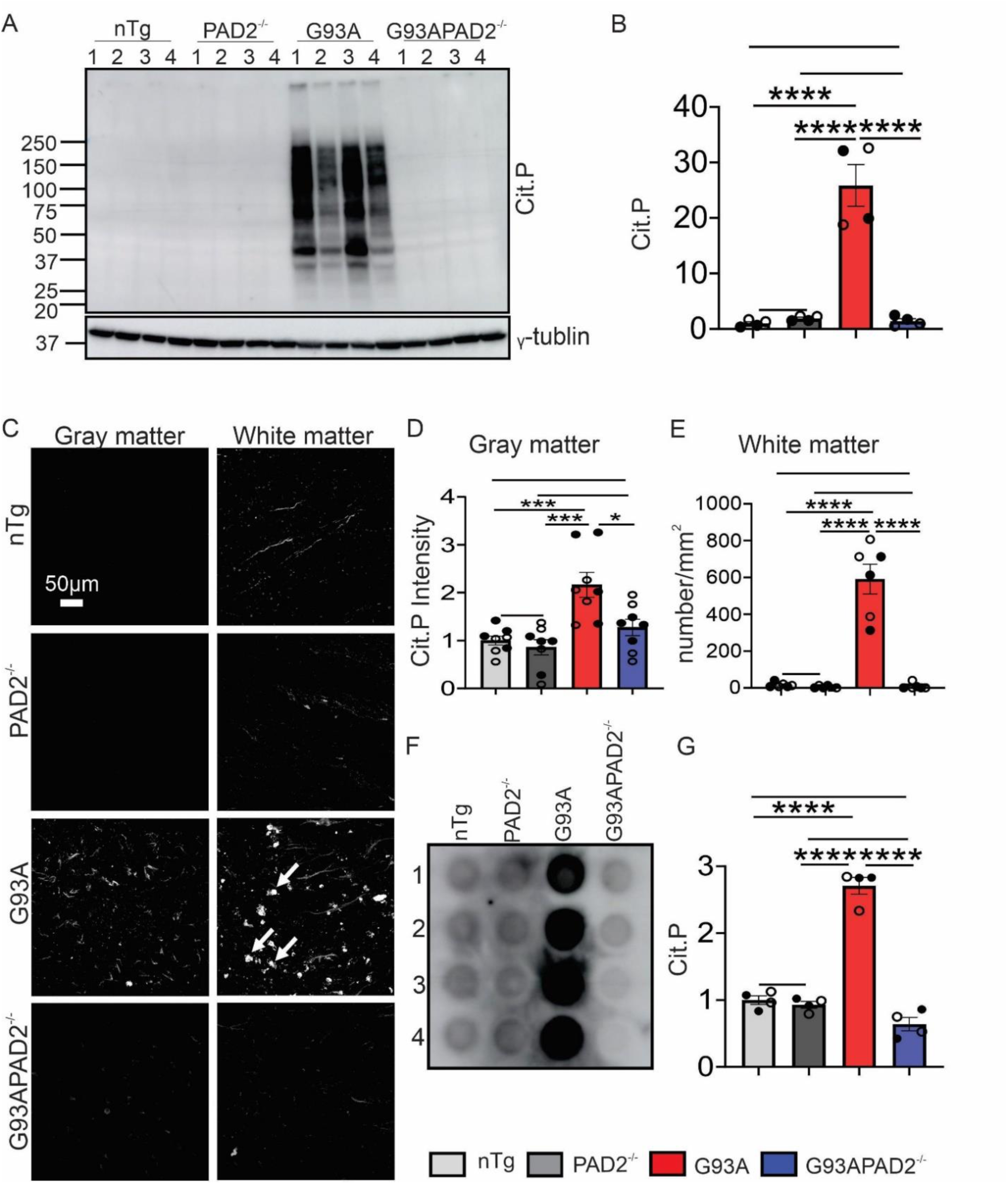
PAD2 knockout blocks the increase of PC (measured by AMC method) in SOD1^G93A^ mice. (A) Western blots for citrullinated proteins in mouse spinal cords. (B) Quantification of lane signal density in blot A. (C) Immunofluorescence staining for citrullinated proteins in ventral spinal cord gray and white matter. Arrows point to examples of citrullinated protein aggregates. (D) Quantification of fluorescent intensity of citrullinated proteins in the spinal cord gray matter. (E) Quantification of citrullinated protein aggregate numbers in the white matter in spinal cord sections as shown C. (F) Filter trap assay for aggregated citrullinated proteins. (G) Quantification of optical density of citrullinated proteins in F. n = 4-8 in each group in all quantifications. The empty and black-filled circles represent the measured values from female and male animals, respectively. Statistics: one-way ANOVA with Bonferroni post-hoc test. *p<0.05. **p<0.01. ***p<0.001. ****p<0.0001. No asterisk indicates p>0.05. The same symbols and statistics are used in the subsequent quantitative graphs unless stated otherwise.

### PAD2 knockout reduced MBP Aggregation in SOD1^G93A^ mice

As we reported previously, the myelin proteins MBP, PLP, and MAG form aggregates (20, 21). Furthermore, the MBP and PLP aggregates are PC-positive whereas MAG aggregates are not. To determine how PC impacts myelin protein aggregation, we performed IF staining for MBP, PLP and MAG in spinal cord white matter. All three proteins showed significant increases in the number of aggregates in SOD1^G93A^ mice compared to the non-transgenic (nTg) controls (Fig. 2A, B). Interestingly, MBP and PLP aggregate numbers were significantly reduced by PAD2KO, whereas MAG aggregates remained unchanged (Fig. 2A, B). To confirm this observation, we performed filter trap assay. PAD2KO significantly attenuated MBP aggregation but the reductions in PLP and MAG were not statistically significant (Fig. 2C, D). These results suggest that PAD2-mediated PC contributes to MBP aggregation, and to a lesser extent, PLP aggregation; but it does not contribute to MAG aggregation.

**Figure 2.**
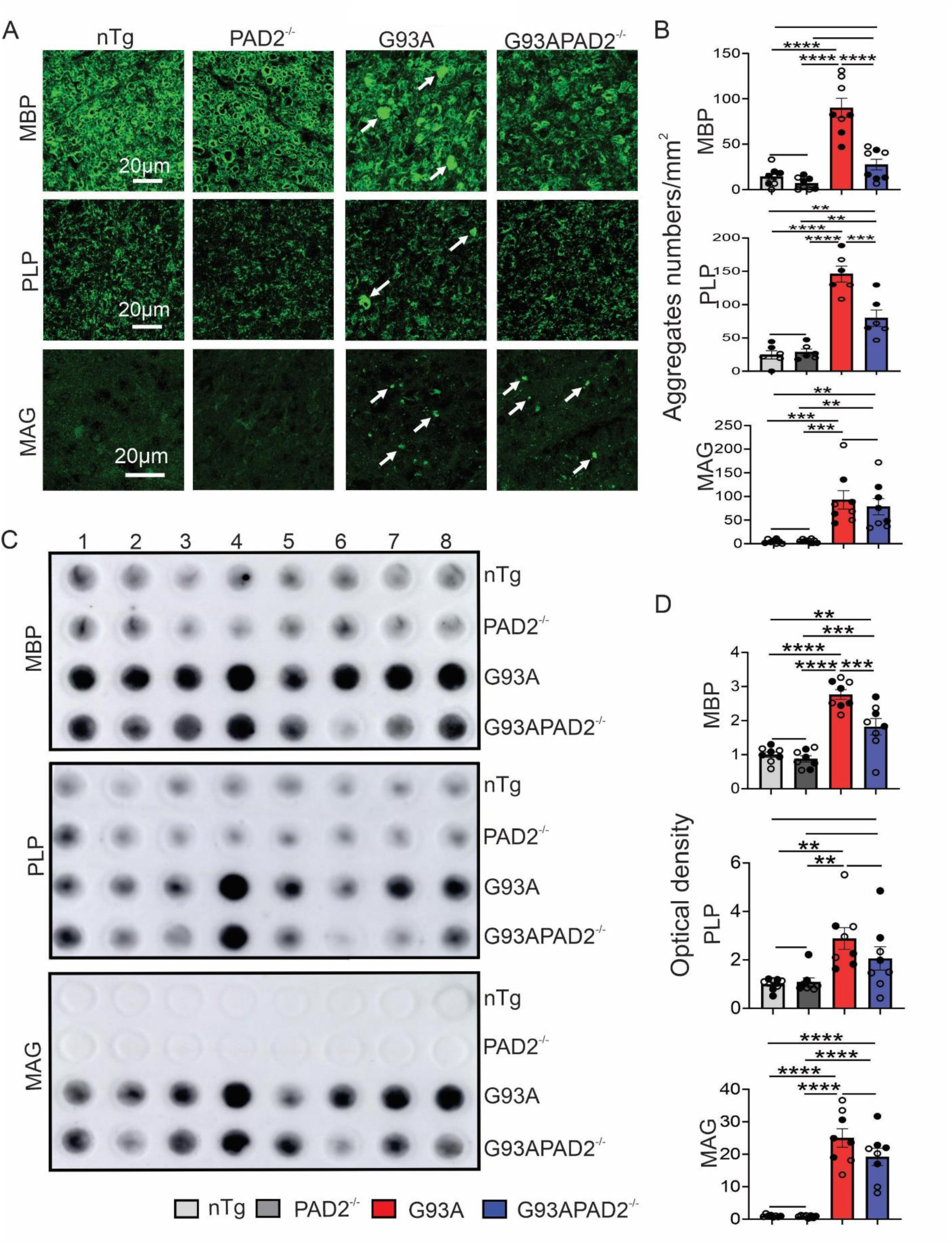
PAD2 knockout reduces MBP aggregation in SOD1^G93A^ mice. (A) Immunofluorescence staining for MBP, PLP, and MAG in spinal cord white matter. Arrows point to protein aggregates. (B) Quantification of MBP, PLP, and MAG aggregates numbers from sections as shown in A. (C) Filter trap assay for MBP, PLP, and MAG protein aggregates in mouse spinal cords. (D) Quantification of optical density of MBP, PLP, and MAG in (C). n = 8 in each group. Symbols and statistics are described in Fig. 1 legend.

Next, we examined whether PAD2KO impacted other well-known protein aggregates such as SOD1 and ubiquitin aggregates. By filter trap assay, we did not find significant changes in SOD1 and ubiquitin aggregates in SOD1^G93A^/PAD2KO mice (Fig. S4), indicating that PAD2-mediated PC does not contribute to mutant SOD1 aggregation.

Could PAD2KO alter myelin gene expression, which in turn changes myelin protein aggregation? To answer this question, we measured the mRNA and protein levels of MBP, MAG, MOG, CNPase and PLP. We did not detect changes in the levels of these molecules (Fig. S5). Another possibility is that PAD2KO impacted mutant SOD1 expression, which might lead to a cascade of downstream beneficial effects. To test this possibility, we also measured mutant SOD1 levels and did not detect changes (Fig. S5). These results show that PAD2 does not impact the expression of myelin genes and mutant SOD1.

### PAD2 knockout enhances motor neuron survival, modulates neuroinflammation and protects myelinated axons

Next, we investigated whether PAD2KO impacted ALS pathology in the spinal cord. By ChAT staining, we found that PAD2KO increased motor neuron survival by ∼20% at the end-stage in SOD1^G93A^ mice (Fig. 3A, B), suggesting that PAD2KO in SOD1^G93A^ enhances neuronal survival. Because the most conspicuous increase in PAD2 and PC is in reactive astrocytes (20, 21), we performed IF staining to evaluate gliosis using antibodies against GFAP (astrocyte), Iba1 (microglia), CD68 (activated microglia), Olig2 (oligodendrocyte), and NG2 (oligodendrocyte progenitor cells). Quantification of fluorescent staining intensities of these markers did not reveal differences caused by PAD2KO in the spinal cord gray and white matters (Fig. S6), with the only exception of Iba1 staining in the white matter, where the IF intensity lowered ∼25% in SOD1^G93A^/PAD2^-/-^ mice compared with SOD1^G93A^ mice (Fig. 3C, D).

**Figure 3.**
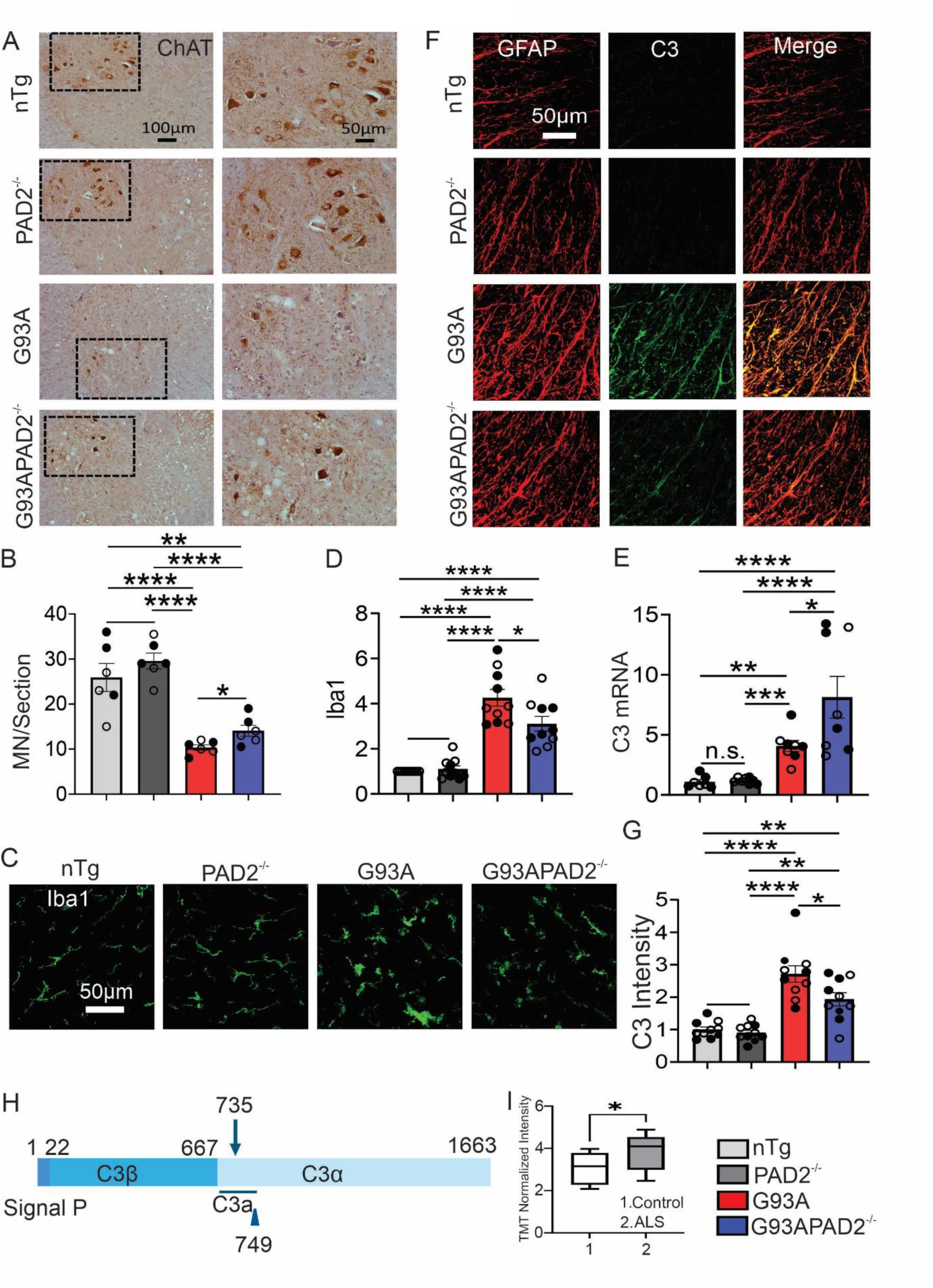
PAD2 knockout enhances neuronal survival, alleviates white matter microgliosis, and reduces C3 protein levels in SOD1^G93A^ mice. (A) Immunohistochemistry staining for ChAT in the spinal cord ventral horn. The right panels are enlarged views of the boxed areas on the left. (B) Quantification of motor neuron numbers in the lumber spinal cord ventral horn in A. (C) Immunofluorescence staining for Iba1 in white matter. (D) Quantification of fluorescent intensity of Iba1 (arbitrary units) in sections as illustrated in C. (E) qPCR quantification of C3 mRNA. (F) Double immunofluorescence staining for GFAP and C3 in white matter. (G) Quantification of fluorescent intensity of C3 staining in sections as illustrated in F. (H) Schematic illustration of C3 protein showing citrullinated site at R735cit. (I) Box plot showing TMT normalized intensities comparing C3 citrullinated peptides from healthy controls and ALS cases. Symbols and statistics for B, D, E, G are described in Fig. 1 legend. n = 6-10 in each group. For I, Student’s t-test was used to compare between control and ALS. n=8 for controls and 9 for ALS.

Given the reduction in Iba1 staining intensity and PC’s involvement in neuroinflammation (25–27), we further examined the inflammation status using a battery of inflammation markers, including: TNF-α, TGF-β1, TGF-β2, IL-6, IL-1β, C1qa, C3, CXCL-1, CX3CL-1, CX3CR1, CCR2, CD86, and CD206 (28–32). Except for IL-6, CX3CL-1, and CCR2, the mRNA levels of all the markers were increased in SOD1^G93A^ mice (Fig. S7). However, only C3 mRNA was further increased by PAD2KO in SOD1^G93A^ mice (Fig. 3E). To determine the C3 protein levels, we performed double IF staining for GFAP and C3, because previous reports showed that C3 expression was increased in astrocytes (33, 34). We confirmed that C3 was increased in reactive astrocytes in SOD1^G93A^ mice (Fig. 3F). Surprisingly, PAD2KO reduced C3 protein levels in astrocytes (Fig 3F, G; Fig. S8A, B), suggesting that citrullination regulates C3 protein levels. To determine whether C3 is citrullinated, we searched the published ALS proteomic datasets using the ionFinder algorithm (24) and found that R735 is citrullinated to a greater extent in samples obtained from the cerebral spinal fluid (CSF) of ALS patients versus healthy controls (23) (Fig. 3H, I; Fig. S8C). Interestingly, the R735W mutation causes atypical hemolytic uremic syndrome (aHUS) presumably due to a gain of function leading to dysregulation of C3 activity (35), thus suggesting that the R735 is a functionally impactful residue.

Considering the reduction of myelin protein aggregates and Iba1 and C3 levels in the white matter, we examined axon integrity by IF staining for MBP and neurofilament light chain (NFL). In the nTg and PAD2KO controls, the staining showed a typical normal pattern, where the central axonal NFL signal was surrounded by myelin represented by the MBP signal (Fig. 4A). By contrast, in SOD1^G93A^ mice, the NFL signal was reduced (Fig. 4A, B), and MBP ring-like structure collapsed, reflecting degeneration of axons (Fig. 4A, arrows and arrowheads). The overall MBP staining intensity did not change from the controls (Fig. 4A, C), likely because the MBP from degenerating axons formed aggregates rather than being cleared. Interestingly, in SOD1^G93A^/PAD2KO mice, there was a noticeable preservation of the pattern of central axonal NFL surrounded by MBP, and the MBP aggregates were absent (Fig. 4A, B). To confirm this observation, we evaluated axon integrity by toluidine blue staining. In the nTg and PAD2KO controls, the circular shaped myelinated axons showed their normal pattern (Fig 4D). However, in the SOD1^G93A^ mice, the normal pattern was disrupted, and irregularly shaped and degenerated myelin structures were numerous (Fig. 4D, arrowheads). In SOD1^G93A^/PAD2KO mice, the myelinated axons were preserved, and the degenerating axon profiles were reduced (Fig. 4D, E). Additional observations by electron microscopy confirmed that PAD2KO preserved axons in the spinal cord (Fig. 4F).

**Figure 4.**
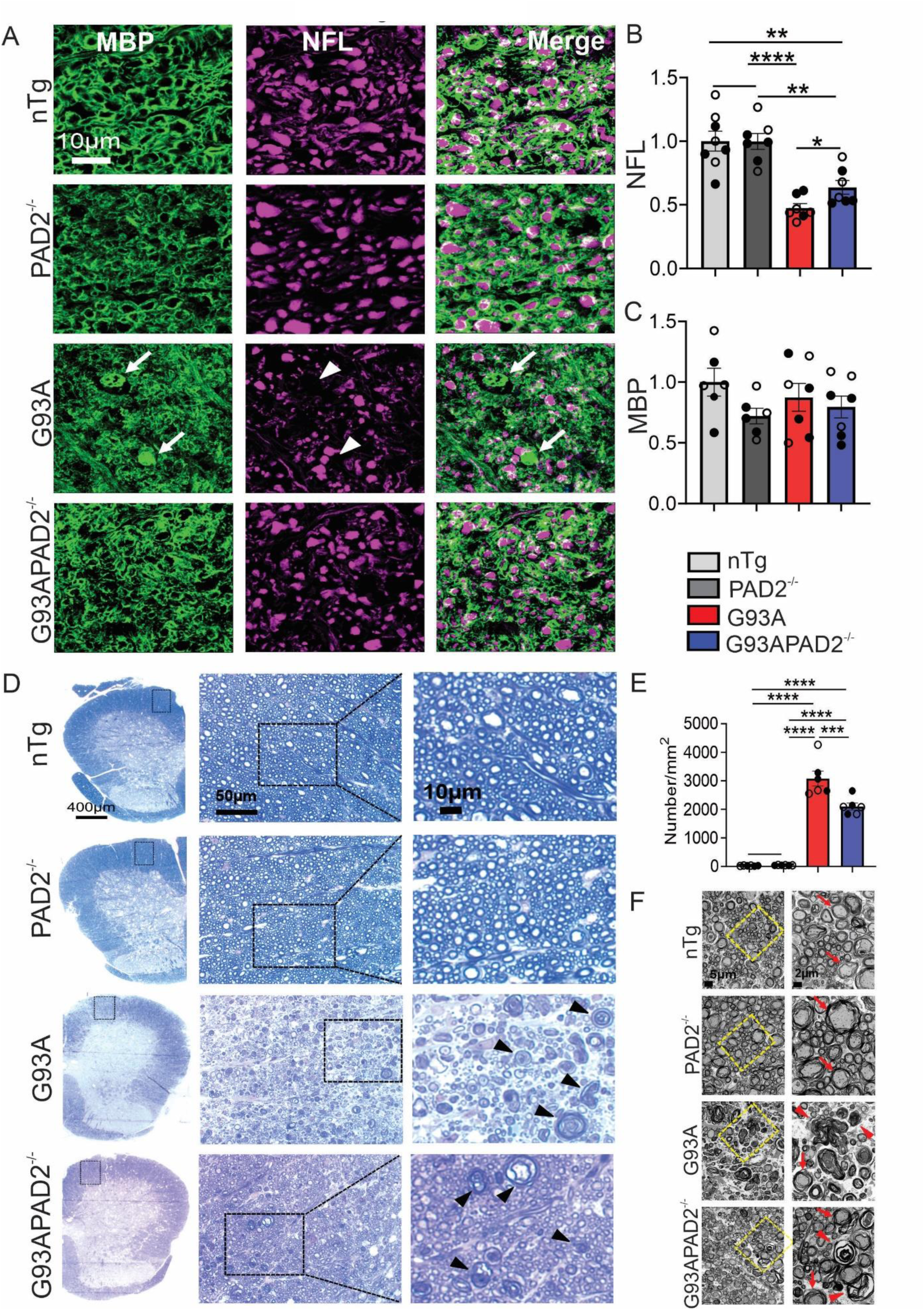
PAD2 knockout attenuated myelin and axonal degeneration in spinal cord white matter of SOD1^G93A^ mice. (A) Double immunofluorescence staining for MBP and NFL in ventral white matter. White arrows point to MBP aggregates. White arrowheads point to areas of axon loss. (B, C) Quantification of fluorescent intensity of NFL and MBP in ventral white matter in A. (D) Toluidine blue stained spinal cord sections. Black arrowheads point to degenerating axons. (E) Quantification of degenerating axons in the ventral spinal cord white matter as shown in D. (F) EM images of ventral white matter. The boxed areas in the left are magnified on the right. Red arrows point to intact axons, and red arrowheads point to degenerating axons. n = 6-8 in each group. Symbols and statistics are described in Fig. 1.

### PAD2 knockout protects motor neuron axons and neuromuscular junctions

To determine whether PAD2KO also protected the motor and sensory axons that project to the periphery, we examined ventral and dorsal roots. In SOD1^G93A^ mice, there was great devastation and loss of ventral and dorsal root axons, leading to a significant reduction in the number of surviving axons compared to the controls (Fig. 5). PAD2KO improved the axon integrity and increased the surviving axons in SOD1^G93A^ mice (Fig. 5).

**Figure 5.**
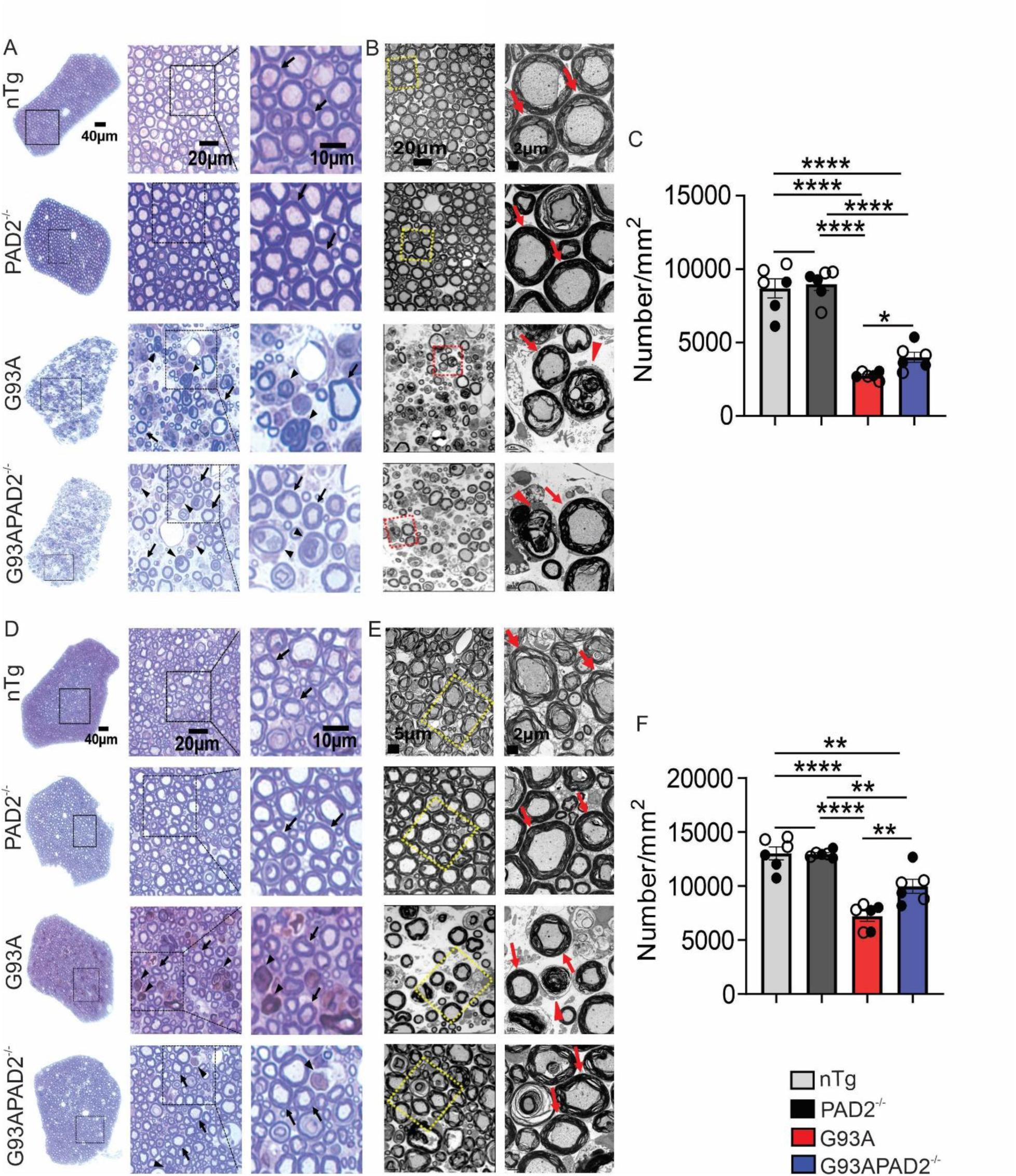
PAD2 knockout protects myelin and axonal integrity in spinal roots of SOD1^G93A^ mice. (A) Toluidine blue staining of L5 ventral root axons. (B) EM images of L5 ventral root axons. (C) Quantification of intact L5 ventral root axons from images as shown in A. (D) Toluidine blue staining of L5 dorsal root axons. (E) TEM of L5 dorsal root axons. (F) Quantification of intact L5 dorsal root axons from images as shown in D. Black and Red arrows point to intact myelinated axons. Black and red arrowheads point to degenerated myelinated axons. n = 6 in all groups. Symbols and statistics are described in Fig. 1.

Next, we investigated whether PAD2KO impacted neuromuscular junctions (NMJs) by triple IF staining using α-Bungarotoxin as a marker of postsynaptic endplate, and synaptophysin and NFL as markers of presynaptic terminals. We quantified NMJ integrity using two methods. First, we quantified NMJ occupancy, which was defined as the percentage of α-Bungarotoxin stained endplate areas that overlapped with the areas stained with presynaptic marker synaptophysin. In nTg and PAD2KO controls, the NMJ occupancy averaged ∼60% (Fig.6A, B). In SOD1^G93A^ mice, NMJ occupancy averaged only ∼10% (Fig. 6A, B). By contrast, NMJ occupancy averaged ∼30% in SOD1^G93A^/PAD2KO mice (Fig. 6A, B), improving from SOD1^G93A^ mice by ∼3 fold. Second, we quantified the NMJ contact, which was defined as the percentage of endplates contacted by the presynaptic terminals. NMJ contact averaged nearly 100% in nTg and PAD2KO controls, but only ∼20% in SOD1^G93A^ mice (Fig. 6A, C). By contrast, in SOD1^G93A^/PAD2KO mice, NMJ contact averaged nearly 60% (Fig. 6A, C), also improving from SOD1^G93A^ mice by ∼3 fold. Thus, PAD2 knockout ameliorates NMJ denervation in SOD1^G93A^ mice.

**Figure 6.**
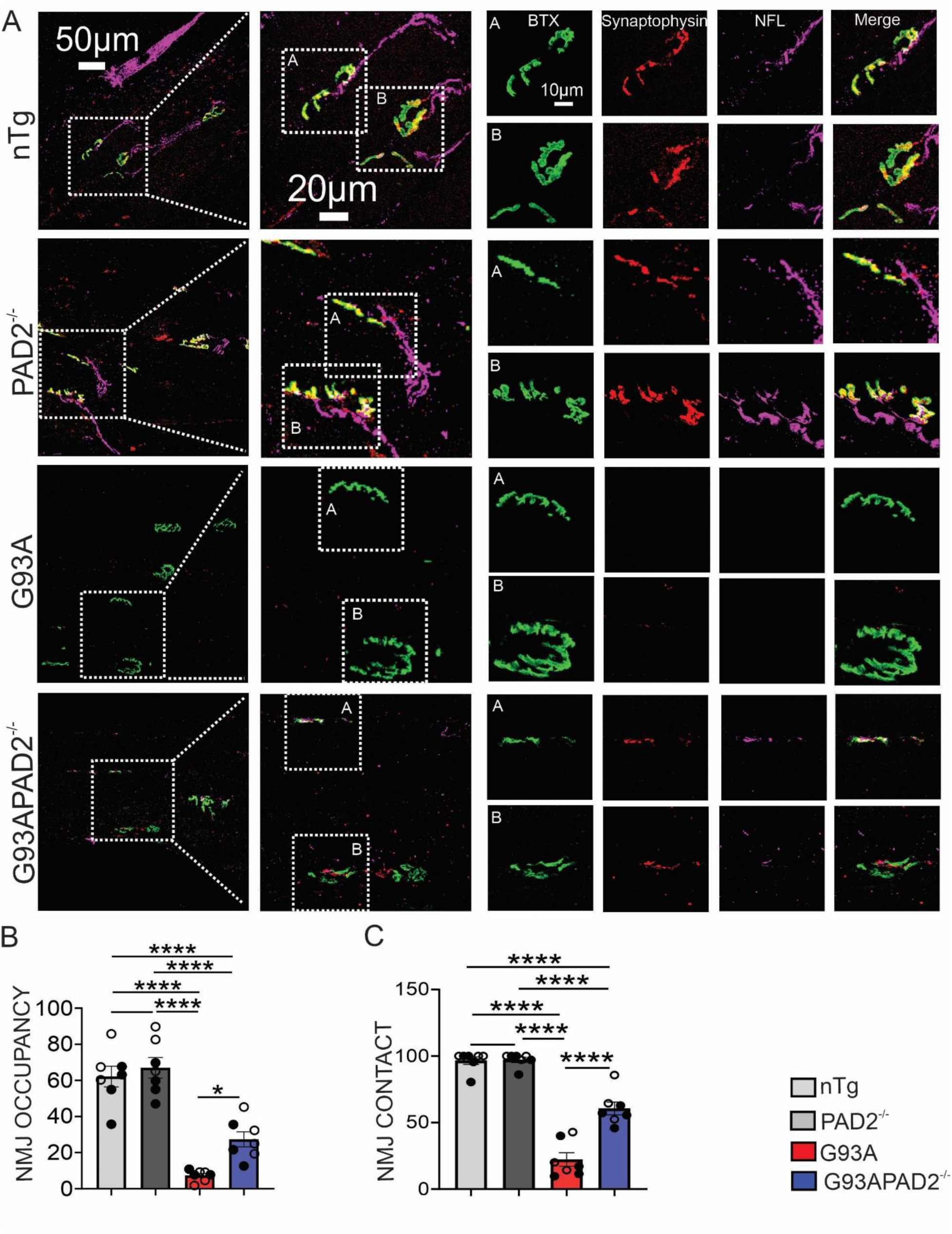
PAD2 knockout protects neuromuscular junction (NMJ) against denervation in SOD1^G93A^ mice. (A) Triple immunofluorescent staining of gastrocnemius muscle sections for α-bungarotoxin (BTX) (green), synaptophysin (red), and NFL (purple). (B) Quantification of postsynaptic NMJ occupancy by the presynaptic terminal, using the formula: NMJ occupancy = (synaptophysin-stained area/α -bungarotoxin-stained area) * 100. (C) Quantification of postsynaptic NMJs contacted by the presynaptic terminal by counting the number of BTX-stained postsynaptic terminals that have an overlap with synaptophysin signal. n = 7 in all groups. Symbols and statistics are described in Fig. 1.

### PAD2 knockout alters the course of disease progression in SOD1^G93A^ mice

To evaluate the overall impact of PAD2KO on the disease phenotype, we monitored the body weight of SOD1^G93A^ and SOD1^G93A^/PAD2KO mice during the disease progression. The average body weight of SOD1^G93A^ mice peaked at age 16 weeks for the males and 18 weeks for the females. By contrast, the average body weight of SOD1^G93A^/PAD2KO mice peaked earlier, at 14 weeks for both males and females (Fig. 7A), suggesting that PAD2KO accelerated disease onset. Analysis of the body weight peaks in individual mice revealed the same trend (Table S4). The median body weight peak of SOD1^G93A^ mice was at 15 weeks for the males and 17 weeks for the females. By contrast, the median body weight peak of SOD1^G93A^/PAD2KO mice occurred earlier, at 14 weeks for both the males and the females (Fig. 7B). We also computed the lifespan and found that PAD2KO significantly extended median survival of male mice by 14 days (Log rank test, p = 0.026) and female mice by 6 days (Log rank test, p = 0.86) (Fig. 7C, Table S4). Overall, PAD2KO accelerated disease onset and slowed disease progression slightly but the differences did not reach statistical significance (Table S4).

**Figure 7.**
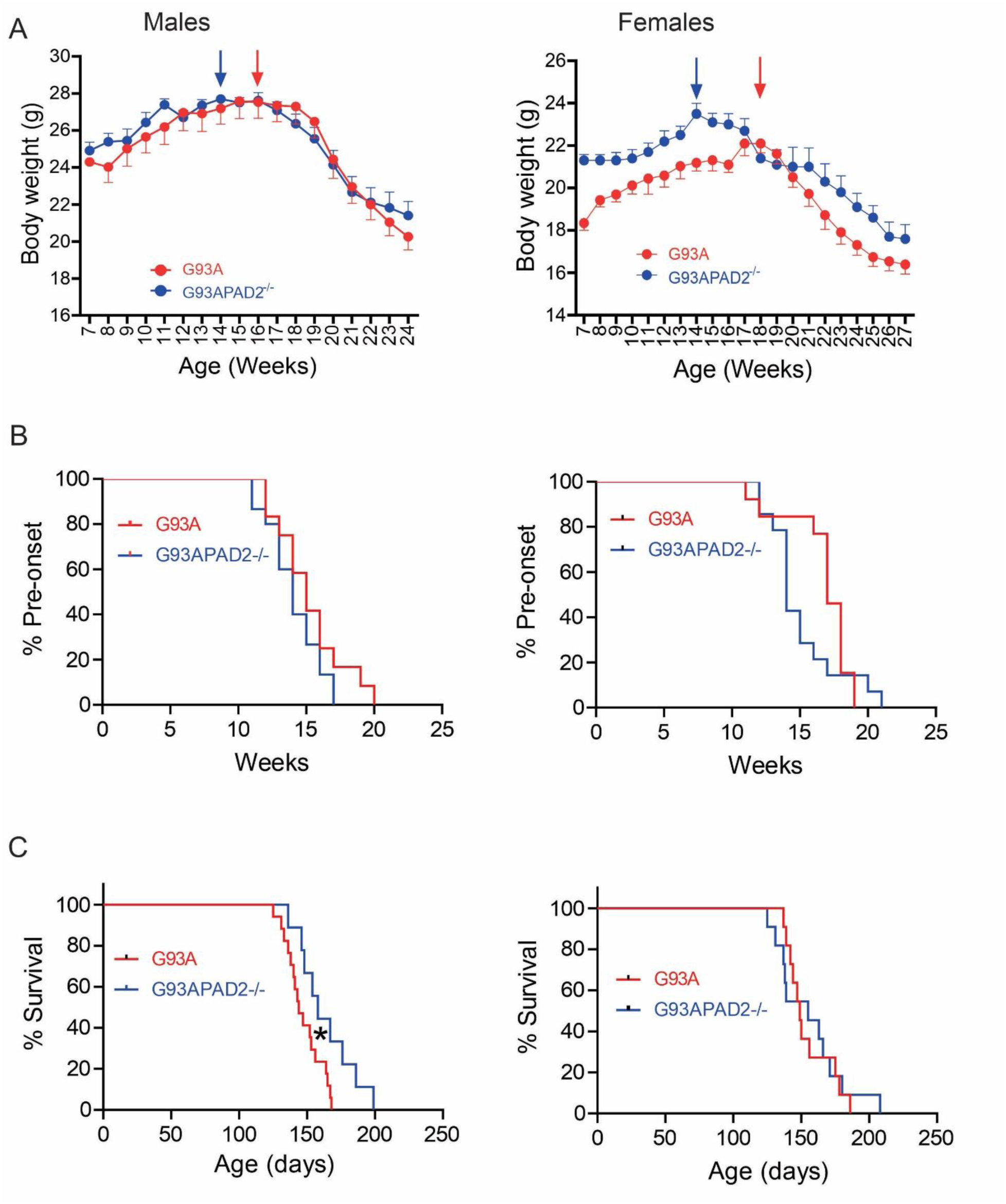
PAD2 knockout modulates the course of disease in SOD1^G93A^ mice. (A) Weekly body weight averaged from 9-15 mice in each data-point. Arrows point to peak weight. (B) Kaplan-Meier Pre-disease onset curves for percent of mice peak body weight in G93A/PAD2^-/-^ and G93A mice (C) Kaplan-Meier survival curves. The onset and lifespan medians and averages and statistics are summarized in Table S4.

## Discussion

In this study, we used SOD1^G93A^/PAD2KO mice to understand the role of PAD2-induced citrullination in ALS. Our results show that PAD2KO completely blocked the bulk increase in PC associated with ALS disease progression and mitigated aggregation of citrullinated proteins as exemplified by MBP. Pathologically, PAD2KO modulated neuroinflammation, protected motor neurons, axons, myelin and NMJ. Clinically, PAD2KO extended survival of male SOD1^G93A^ mice and might have modulated disease course by accelerating disease onset but slowing disease progression, although the latter effect did not reach statistical significance. Collectively, these results demonstrate that PAD2 is responsible for the dramatic increase in PC in ALS and PAD2-induced PC contributes to the disease.

In our previous studies, we showed that PAD2 and PC are increased in astrocytes in ALS mouse models and human postmortem tissues (20, 21). Therefore, we hypothesized that PAD2 was responsible for the dramatic increase in PC observed in astrocytes during ALS disease progression. This study provides evidence substantiating this hypothesis, as PAD2KO completely prevented the PC increase in astrocytes (Fig. 1). A surprising finding is that PAD2KO also eliminated citrullinated protein aggregates in the white matter (Fig. 1), indicating that PAD2 is responsible for citrullinated protein aggregates despite the fact that PAD2 did not colocalize with the PC aggregates (20). This finding raises the question as to whether PC causes protein aggregation. Although a definitive answer can only be attained from further studies, the observation that PAD2KO reduced MBP and possibly PLP aggregates (Fig. 2) strongly suggests that PC promotes aggregation of the myelin proteins.

Do other PADs contribute to the increased PC in ALS? Our results do not support such a role. Although low levels of residual PC could be detected in PAD2KO mice, PAD2KO completely abolished the robust increase of PC in the ALS mouse model. Additionally, PAD1 was not detectable and PAD3 and PAD4 were detected but there were no compensatory changes (Fig. S2). Nevertheless, these other PADs are probably responsible for the residual PC in the CNS of the PAD2KO mice. Indeed, several studies reported that PAD4 is expressed in neurons and astrocytes in normal and pathological conditions (36–39). In our study, we attempted to replicate those results and to determine whether PAD2KO alters the patterns of expression of the other PADs. However, our attempt was halted due to the lack of specific PAD4 antibodies (Fig. S3). Alarmingly, of the four PAD4 antibodies that we tested, three also recognized PAD2 strongly and all recognized multiple non-PAD4 bands on Western blots of mouse CNS protein lysates (Fig. S3). Therefore, we caution the interpretation of the previous literature unless the specificity of the PAD4 antibodies were verified with the CNS protein extracts from the species under the study.

The evidence presented in this study suggests that the increased PAD2 activity during ALS disease progression on balance exacerbates the disease. Histological analysis revealed that PAD2KO reduced myelin protein aggregation (Fig. 2), ameliorated microgliosis in the white matter and reduced C3 protein levels (Fig. 3), and enhanced motor neuron survival with improved myelin, axon and NMJ integrity in SOD1^G93A^ mice (Figs. 3-6). Clinically, the effects of PAD2KO are mixed. Compared with SOD1^G93A^ mice, SOD1^G93A^/PAD2KO mice showed a trend of earlier disease onset, but the lifespan was unchanged in females and longer in males (Fig. 7). These changes led to a longer disease duration suggesting a slower disease progression after the onset. These results show the complexity in the role of PAD2 and PC in ALS. Our previous study showed that PAD2 and PC are decreased in motor neurons at the disease onset and thereafter during the disease progression. On the other hand, PAD2 and PC increase in astrocytes as the disease progresses after onset (20). We hypothesize that the reduced PC in neurons weakens neuron health in early disease, and this condition is exacerbated by PAD2KO hence earlier disease onset; on the other hand, the increased PAD2 and PC in astrocytes contributes to neuroinflammation that is blunted by PAD2KO hence slower disease progression. Therefore, the beneficial and detrimental effects of PAD2KO cancel each other, leading to diminished clinical impact. This interpretation is supported by the fact that a large number of proteins are citrullinated in CNS in the SOD1^G93A^ model (20). Some of the PC events may be beneficial while others may be detrimental, thereby leading to mixed clinical outcomes by PAD2KO.

The association of PC with inflammation has been well established (25, 40). Neuroinflammation also contributes to ALS (41). Our results, however, draw a nuanced picture regarding the relationship between PC and neuroinflammation. We measured the expression of numerous inflammation markers and found that most remained unchanged after PAD2KO (Fig. S6, S7). However, microgliosis and C3 complement protein levels were reduced (Fig. 3). C3 is highly expressed in reactive astrocytes and thought to promote neurodegeneration in various neuroinflammatory conditions (42, 43). Indeed, overexpression of C3 in astrocytes increases neuronal loss in PD, whereas decreasing C3 production alleviates neuronal loss in an AD mouse model (44, 45). Our study shows that C3 is citrullinated at R735 in ALS patient CSF (Fig. 3H-J). The R735W mutation causes aHUS, a disease linked to dysregulated hyperactivity of complements (35). The R735Cit could have a similar functional impact on C3 activity and contribute to neuroinflammation. Therefore, the lowered levels of C3 protein in astrocytes could contribute to the beneficial effects of PAD2KO, including protection of motor neurons, myelinated axons and NMJs.

However, the impact of C3 and other complement components on ALS is complex. Lobsiger and coworkers reported that C3 knockout in SOD1^G93A^ mice did not impact on the ALS phenotype (46). Knockouts or inhibition of other complements showed mixed outcomes with some showing benefits while others showing no impact (46–48). Interestingly, Lobsiger and colleagues observed a modest male-specific shortening of survival in SOD1^G37R^/C1q^-/-^ mice, suggesting a modest beneficial effect of C1q induction in males (46). Our observation in SOD1^G93A^/PAD2KO mice, where a significant survival extension was only observed in males (Fig. 7C), similarly suggest a male-specific effect of PC, possibly via modulation of C3 levels and activity. Sex-dimorphic responses to genetic or pharmacological interventions in mutant SOD1 mouse model are not unprecedented. For example, Lim and coworkers reported that heterozygous deletion of the leptin gene only extended the lifespan of the female SOD1^G93A^ mice (49). Conversely, Colognesi and colleagues reported that treatment using REL-1017, an NMDA receptor antagonist, extended survival only in males (50). These observations may not be surprising in the context of literature documenting that ALS develop and progress differently in males and females. ALS incidence and prevalence are about 1.3 times higher in men than in women, and ALS is diagnosed earlier in men than in women (51). Furthermore, male survival is shorter than women after diagnosis (52). Similarly, the lifespan of males is shorter than the females in SOD1^G93A^ mice (53). The mechanisms whereby sex impact ALS phenotype and responses to treatments remain unknown.

## Conclusion

In summary, PAD2KO in SOD1^G93A^ mice blocked the increase of PC and reduced aggregation of myelin proteins. At the cellular level, PAD2KO improved motor neuron survival and myelin, axon and NMJ integrity, and reduced complement C3 protein levels and microgliosis in the white matter in SOD1^G93A^ mice. However, the modulation of clinical phenotype was modest, with only significant extension of survival in male mice and a trend of accelerated onset but slowed progression. These modest clinical phenotype modulations may be a consequence of complex roles of citrullination in different proteins and cell types, with some citrullination events beneficial and others detrimental. The ubiquitous knockout of PAD2 leads to cancellation between the beneficial and detrimental effects. Thus, future studies need to focus on discerning the impact of citrullination on individual proteins and specific cell types.

## Supporting information

Supplemental Table 1: List of qPCR Primers.

Supplemental Table 2: List of Primary Antibodies.

Supplemental Table 3: List of Secondary Antibodies.

Supplemental Figure 1: Genetic knockout of PAD2 eliminates PAD2 expression.

Supplemental Figure 2: PAD2 knockout does not alter other expressions of other PADs.

Supplemental Figure 3: Current commercial PAD4 antibodies are not specific for PAD4 in mouse spinal cord lysate. PAD4 antibodies, including.

Supplemental Figure 4: PAD2 does not contribute to SOD1 and ubiquitin aggregation in SOD1G93A mice.

Supplemental Figure 5: PAD2 knockout does not alter myelin gene expression in SOD1G93A mice.

Supplemental Figure 6: PAD2 knockout does not impact gliosis in the spinal cord of SOD1G93A mice.

Supplemental Figure 7: PAD2 knockout does not impact the expression of many inflammatory markers in the spinal cord of SOD1G93A mice.

Supplemental Figure 8: PAD2 knockout reduces C3 in astrocytes in the spina cord gray matter in SOD1G93A mice.

Supplemental Table 4: The onset and lifespan medians, averages and statistics

Supplemental Table 5: Onset and survival data from individual mice

## Abbreviations

PADs: Protein arginine deiminases
PAD1: Protein arginine deiminase 1
PAD2: Protein arginine deiminase 2
PAD3: Protein arginine deiminase 3
PAD4: Protein arginine deiminase 4
ALS: Amyotrophic lateral sclerosis
SOD1: Superoxide dismutase 1
AMC: Antimodified citrulline
ChAT: Choline acetyl transferase
PC: Protein citrullination
BTX: Bungarotoxin
PAD2KO: Protein arginine deiminase 2 knockout
GFAP: Glial fibrillary acidic protein
MBP: Myelin basic protein
PLP: Proteolipid protein
MAG: Myelin associated glycoprotein
MOG: Myelin oligodendrocyte glycoprotein
CNPase: 2’,3’-cyclic nucleotide 3’-phosphodiesterase
Iba1: Ionized calcium-binding adapter molecule 1
Cd68: Cluster of Differentiation 68
NFL: Neurofilament light chain
C3: Complement C3
Olig2: Oligodendrocyte Transcription Factor 2
Ng2: Neuron-Glial Antigen 2
NMJ: Neuromuscular junction
TMT: Tandem Mass Tag
PTM: Posttranslational modification
AD: Alzheimer’s disease
PD: Parkinson’s disease
MS: Multiple sclerosis
TEM: Transmission electron microscopy
qPCR: Quantitative Polymerase Chain Reaction
TNF-α: Tumor necrosis factor-alpha
TGF-β1: Transforming growth factor beta 1
TGF-β2: Transforming Growth Factor-beta 2
IL-6: Interleukin-6
IL-1β: Interleukin-1 beta
C1qa: Complement C1q A chain
CXCL-1: C-X-C motif chemokine ligand 1
CX3CL-1: C-X3-C motif chemokine ligand 1
CX3CR1: C-X3-C motif chemokine receptor 1
CCR2: C-C motif chemokine receptor 2
CD86: Cluster of Differentiation 86
CD206: Cluster of Differentiation 206
aHUS: Atypical Hemolytic Uremic Syndrome

## Acknowledgements

The authors are grateful for the support from the core facilities at the University of Massachusetts Chan Medical School, including Morphology Core, Confocal Microscopy Core managed by Dr. Kerstin Nundel, Electron Microscopy Core managed by Dr. Gregory Hendricks and Keith Reddig, and the Department of Animal Medicine. We thank Dr. Scott Coonrod (Cornell University, Ithaca, New York, USA) for generously providing us with the PAD2KO mice.

## Funding

This work was supported by grants from the Angel Fund for ALS Research, NIH/NINDS RO1-NS118145 to ZX and PRT, and NIH/NIGMS R35 GM118112 to PRT

## CRediT authorship contribution statement

Issa O. Yusuf: Writing – review & editing, Writing – original draft, Visualization, Validation, Resources, Project administration, Methodology, Investigation, Formal analysis, Data curation, Conceptualization.

Rogerio L.A. Silva: Resources, Investigation, Methodology.

George Amoako: Resources, Investigation

Paul R. Thompson: Writing – review & editing, Supervision, Funding acquisition, Conceptualization.

Zuoshang Xu: Writing – review & editing, Writing – original draft, Supervision, Resources, Project administration, Methodology, Investigation, Funding acquisition, Formal analysis, Data curation, Conceptualization.

## Declaration of competing interest

PRT holds equity in Padlock Therapeutics a subsidiary of Bristol Myers Squibb. The other authors have no competing interests to declare that are relevant to this article.

## Data availability

All data supporting the conclusions of this article are included within the article and its supplementary materials.

## References

1. Yuan L, Yang Y, Guo Y, Deng H. Genetic architecture of amyotrophic lateral sclerosis: a comprehensive review. J Genet Genomics. 2025;52(10):1155–76.

2. Zhou T, Ahmad TK, Gozda K, Truong J, Kong J, Namaka M. Implications of white matter damage in amyotrophic lateral sclerosis (Review). Mol Med Rep. 2017;16(4):4379–92.

3. Taylor JP, Brown RH, Jr., Cleveland DW. Decoding ALS: from genes to mechanism. Nature. 2016;539(7628):197–206.

4. Quigley SE, Quigg KH, Goutman SA. Genetic and Mechanistic Insights Inform Amyotrophic Lateral Sclerosis Treatment and Symptomatic Management: Current and Emerging Therapeutics and Clinical Trial Design Considerations. CNS Drugs. 2025;39(10):949–93.

5. Shimizu M, Okuno T. Disruption of neuronal actin barrier promotes the entry of disease-implicated proteins to exacerbate amyotrophic lateral sclerosis pathology. Neural Regen Res. 2025;20(9):2589–90.

6. Mondal S, Thompson PR. Protein Arginine Deiminases (PADs): Biochemistry and Chemical Biology of Protein Citrullination. Acc Chem Res. 2019;52(3):818–32.

7. Fujisaki M, Sugawara K. Properties of peptidylarginine deiminase from the epidermis of newborn rats. J Biochem. 1981;89(1):257–63.

8. Jang B, Jin JK, Jeon YC, Cho HJ, Ishigami A, Choi KC, et al. Involvement of peptidylarginine deiminase-mediated post-translational citrullination in pathogenesis of sporadic Creutzfeldt-Jakob disease. Acta Neuropathol. 2010;119(2):199–210.

9. Lange S, Rocha-Ferreira E, Thei L, Mawjee P, Bennett K, Thompson PR, et al. Peptidylarginine deiminases: novel drug targets for prevention of neuronal damage following hypoxic ischemic insult (HI) in neonates. J Neurochem. 2014;130(4):555–62.

10. Lazarus RC, Buonora JE, Flora MN, Freedy JG, Holstein GR, Martinelli GP, et al. Protein Citrullination: A Proposed Mechanism for Pathology in Traumatic Brain Injury. Front Neurol. 2015;6:204.

11. Mastronardi FG, Wood DD, Mei J, Raijmakers R, Tseveleki V, Dosch HM, et al. Increased citrullination of histone H3 in multiple sclerosis brain and animal models of demyelination: a role for tumor necrosis factor-induced peptidylarginine deiminase 4 translocation. J Neurosci. 2006;26(44):11387–96.

12. Moscarello MA, Pritzker L, Mastronardi FG, Wood DD. Peptidylarginine deiminase: a candidate factor in demyelinating disease. J Neurochem. 2002;81(2):335–43.

13. Nicholas AP. Dual immunofluorescence study of citrullinated proteins in Parkinson diseased substantia nigra. Neurosci Lett. 2011;495(1):26–9.

14. Wu Z, Deng Q, Pan B, Alam HB, Tian Y, Bhatti UF, et al. Inhibition of PAD2 Improves Survival in a Mouse Model of Lethal LPS-Induced Endotoxic Shock. Inflammation. 2020;43(4):1436–45.

15. Dong T, Barasa L, Yu X, Ouyang W, Shao L, Quan C, et al. AFM41a: A Novel PAD2 Inhibitor for Sepsis Treatment-Efficacy and Mechanism. Int J Biol Sci. 2024;20(13):5043–55.

16. Yasuda H, Saito M, Hayashi S, Kato S. Peptidylarginine deiminase 2 contributes to pathogenesis in trinitrobenzenesulfonic acid-induced colitis through macrophage extracellular trap-independent pathways. Sci Rep. 2025;15(1):40583.

17. Zhou J, Biesterveld BE, Li Y, Wu Z, Tian Y, Williams AM, et al. Peptidylarginine Deiminase 2 Knockout Improves Survival in hemorrhagic shock. Shock. 2020;54(4):458–63.

18. Tian Y, Qu S, Alam HB, Williams AM, Wu Z, Deng Q, et al. Peptidylarginine deiminase 2 has potential as both a biomarker and therapeutic target of sepsis. JCI Insight. 2020;5(20).

19. Dawood ZS, Liggett MR, Wang B, Zhang K, Jin G, Ouyang W, et al. Role of peptidylarginine deiminase 2 in a murine model of traumatic brain injury. J Trauma Acute Care Surg. 2026;100(2):189–97.

20. Yusuf IO, Qiao T, Parsi S, Tilvawala R, Thompson PR, Xu Z. Protein citrullination marks myelin protein aggregation and disease progression in mouse ALS models. Acta Neuropathol Commun. 2022;10(1):135.

21. Yusuf IO, Parsi S, Ostrow LW, Brown RH, Thompson PR, Xu Z. PAD2 dysregulation and aberrant protein citrullination feature prominently in reactive astrogliosis and myelin protein aggregation in sporadic ALS. Neurobiol Dis. 2024;192:106414.

22. Asaga H, Senshu T. Combined biochemical and immunocytochemical analyses of postmortem protein deimination in the rat spinal cord. Cell Biol Int. 1993;17(5):525–32.

23. Trautwig AN, Fox EJ, Dammer EB, Shantaraman A, Ping L, Duong DM, et al. Network analysis of the cerebrospinal fluid proteome reveals shared and unique differences between sporadic and familial forms of amyotrophic lateral sclerosis. Mol Neurodegener. 2025;20(1):58.

24. Maurais AJ, Salinger AJ, Tobin M, Shaffer SA, Weerapana E, Thompson PR. A Streamlined Data Analysis Pipeline for the Identification of Sites of Citrullination. Biochemistry. 2021;60(38):2902–14.

25. Yusuf IO, Camille W, Thompson PR, Xu Z. Protein Citrullination in Amyotrophic Lateral Sclerosis and Other Neurodegenerative Diseases. J Exp Neurol. 2024;5(4):183–91.

26. Femiano C, Bruno A, Gilio L, Buttari F, Dolcetti E, Galifi G, et al. Inflammatory signature in amyotrophic lateral sclerosis predicting disease progression. Sci Rep. 2024;14(1):19796.

27. Petrozziello T, Mills AN, Vaine CA, Penney EB, Fernandez-Cerado C, Legarda GPA, et al. Neuroinflammation and histone H3 citrullination are increased in X-linked Dystonia Parkinsonism post-mortem prefrontal cortex. Neurobiol Dis. 2020;144:105032.

28. Endo F, Komine O, Fujimori-Tonou N, Katsuno M, Jin S, Watanabe S, et al. Astrocyte-derived TGF-beta1 accelerates disease progression in ALS mice by interfering with the neuroprotective functions of microglia and T cells. Cell Rep. 2015;11(4):592–604.

29. Jensen BK, McAvoy KJ, Heinsinger NM, Lepore AC, Ilieva H, Haeusler AR, et al. Targeting TNFalpha produced by astrocytes expressing amyotrophic lateral sclerosis-linked mutant fused in sarcoma prevents neurodegeneration and motor dysfunction in mice. Glia. 2022;70(7):1426–49.

30. Lee JD, Levin SC, Willis EF, Li R, Woodruff TM, Noakes PG. Complement components are upregulated and correlate with disease progression in the TDP-43(Q331K) mouse model of amyotrophic lateral sclerosis. J Neuroinflammation. 2018;15(1):171.

31. Xu CZ, Huan X, Luo SS, Zhong HH, Zhao CB, Chen Y, et al. Serum cytokines profile changes in amyotrophic lateral sclerosis. Heliyon. 2024;10(7):e28553.

32. Zhao W, Beers DR, Hooten KG, Sieglaff DH, Zhang A, Kalyana-Sundaram S, et al. Characterization of Gene Expression Phenotype in Amyotrophic Lateral Sclerosis Monocytes. JAMA Neurol. 2017;74(6):677–85.

33. Guttenplan KA, Weigel MK, Adler DI, Couthouis J, Liddelow SA, Gitler AD, et al. Knockout of reactive astrocyte activating factors slows disease progression in an ALS mouse model. Nat Commun. 2020;11(1):3753.

34. Taha DM, Clarke BE, Hall CE, Tyzack GE, Ziff OJ, Greensmith L, et al. Astrocytes display cell autonomous and diverse early reactive states in familial amyotrophic lateral sclerosis. Brain. 2022;145(2):481–9.

35. Fremeaux-Bacchi V, Miller EC, Liszewski MK, Strain L, Blouin J, Brown AL, et al. Mutations in complement C3 predispose to development of atypical hemolytic uremic syndrome. Blood. 2008;112(13):4948–52.

36. Acharya NK, Nagele EP, Han M, Coretti NJ, DeMarshall C, Kosciuk MC, et al. Neuronal PAD4 expression and protein citrullination: possible role in production of autoantibodies associated with neurodegenerative disease. J Autoimmun. 2012;38(4):369–80.

37. Seol SI, Oh SA, Davaanyam D, Lee JK. Blocking peptidyl arginine deiminase 4 confers neuroprotective effect in the post-ischemic brain through both NETosis-dependent and - independent mechanisms. Acta Neuropathol Commun. 2025;13(1):33.

38. Navneet S, Wilson K, Rohrer B. Muller Glial Cells in the Macula: Their Activation and Cell-Cell Interactions in Age-Related Macular Degeneration. Invest Ophthalmol Vis Sci. 2024;65(2):42.

39. Wizeman JW, Mohan R. Expression of peptidylarginine deiminase 4 in an alkali injury model of retinal gliosis. Biochem Biophys Res Commun. 2017;487(1):134–9.

40. Ciesielski O, Biesiekierska M, Panthu B, Soszynski M, Pirola L, Balcerczyk A. Citrullination in the pathology of inflammatory and autoimmune disorders: recent advances and future perspectives. Cell Mol Life Sci. 2022;79(2):94.

41. Mead RJ, Shan N, Reiser HJ, Marshall F, Shaw PJ. Amyotrophic lateral sclerosis: a neurodegenerative disorder poised for successful therapeutic translation. Nat Rev Drug Discov. 2023;22(3):185–212.

42. Muhamad NA, Masutani K, Furukawa S, Yuri S, Toriyama M, Matsumoto C, et al. Astrocyte-Specific Inhibition of the Primary Cilium Suppresses C3 Expression in Reactive Astrocyte. Cell Mol Neurobiol. 2024;44(1):48.

43. Zhang H, Jin Q, Li J, Wang J, Li M, Yin Q, et al. Astrocyte-derived complement C3 facilitated microglial phagocytosis of synapses in Staphylococcus aureus-associated neurocognitive deficits. PLoS Pathog. 2025;21(4):e1013126.

44. Chi X, Yin S, Sun Y, Kou L, Zou W, Wang Y, et al. Astrocyte-neuron communication through the complement C3-C3aR pathway in Parkinson’s disease. Brain Behav Immun. 2025;123:229–43.

45. Shi Q, Chowdhury S, Ma R, Le KX, Hong S, Caldarone BJ, et al. Complement C3 deficiency protects against neurodegeneration in aged plaque-rich APP/PS1 mice. Sci Transl Med. 2017;9(392).

46. Lobsiger CS, Boillee S, Pozniak C, Khan AM, McAlonis-Downes M, Lewcock JW, et al. C1q induction and global complement pathway activation do not contribute to ALS toxicity in mutant SOD1 mice. Proc Natl Acad Sci U S A. 2013;110(46):E4385–92.

47. Woodruff TM, Lee JD, Noakes PG. Role for terminal complement activation in amyotrophic lateral sclerosis disease progression. Proc Natl Acad Sci U S A. 2014;111(1):E3–4.

48. Woodard GE. C3 and C5 complement cascade activation in brain injury and disease: Molecular mechanisms, pathological roles, and therapeutic implications. Neurochem Int. 2026;196:106161.

49. Lim MA, Bence KK, Sandesara I, Andreux P, Auwerx J, Ishibashi J, et al. Genetically altering organismal metabolism by leptin-deficiency benefits a mouse model of amyotrophic lateral sclerosis. Hum Mol Genet. 2014;23(18):4995–5008.

50. Colognesi M, Shkodra A, Gabbia D, Kawamata H, Manfredi PL, Manfredi G, et al. Sex-dependent effects of the uncompetitive N-methyl-D-aspartate receptor antagonist REL-1017 in G93A-SOD1 amyotrophic lateral sclerosis mice. Front Neurol. 2024;15:1384829.

51. Renzini A, Pigna E, Rocchi M, Cedola A, Gigli G, Moresi V, et al. Sex and HDAC4 Differently Affect the Pathophysiology of Amyotrophic Lateral Sclerosis in SOD1-G93A Mice. Int J Mol Sci. 2022;24(1).

52. Grassano M, Moglia C, Palumbo F, Koumantakis E, Cugnasco P, Callegaro S, et al. Sex Differences in Amyotrophic Lateral Sclerosis Survival and Progression: A Multidimensional Analysis. Ann Neurol. 2024;96(1):159–69.

53. Pfohl SR, Halicek MT, Mitchell CS. Characterization of the Contribution of Genetic Background and Gender to Disease Progression in the SOD1 G93A Mouse Model of Amyotrophic Lateral Sclerosis: A Meta-Analysis. J Neuromuscul Dis. 2015;2(2):137–50.

