## Supplemental Table 1: List of qPCR Primers. for "PAD2 knockout reduces myelin protein aggregates, modulates neuroinflammation and protects motor neurons, axons and neuromuscular junction in a SOD1-ALS mouse model"

**Table S1: List of qPCR Primers.**

| Gene name | Forward primer | Reverse primer |
|---|---|---|
| PAD1 | CTTCAAGGTGAAGGTGTCATACT | CGACATCAAGGGACACATCAA |
| PAD2 | GTTATGTTCAAGGGCCTGGGAGGCATG | TAGCACGATCATGTTCACCATGTTAGG |
| PAD3 | TTCTCCGAGACCCCCATCTT | TTATTCCTCACCCGGCACAC |
| PAD4 | TCTTTGTGGGTCACGTGGATGAGT | AGCTCCTGGAACAGCTGATAGCAA |
| RPL32 | ATGGCTCCTTCGTTGCTG C | CTGGACGGCTAATGCTGGT |
| C3 | CGCAACGAACAGGTGGAGATCA | CTGGAAGTAGCGATTCTTGGCG |
| PLP | CCAGAATGTATGGTGTTCTCCC | GGCCCATGAGTTTAAGGACG |
| MBP | GACCATCCAAGAAGACCCCAC | GCCATAATGGGTAGTTCTCGTGT |
| MAG | CTGCCGCTGTTTTGGATAATGA | CATCGGGGAAGTCGAAACGG |
| MOG | AGCTGCTTCCTCTCCCTTCTC | ACTAAAGCCCGGATGGGATAC |
| CNPase | GCAGGAGGTGGTGAAGAGAT | CAGATGGCTTGTCCAGATCA |
| C1qa | GTGGCTGAAGATGTCTGCCGAG | TTAAAACCTCGGATACCAGTCCG |
| CD86 | ACGATGGACCCCAGATGCACCA | GCGTCTCCACGGAAACAGCA |
| CD206 | TCAGCTATTGGACGCGAGGCA | TCCGGGTTGCAAGTTGCCGT |
| IL-6 | AAC GATGATGCACTTGCAGA | GAGCATTGGAAATTGGGGTA |
| TGF-β1 | TGATACGCCTGAGTGGCTGTCT | CACAAGAGCAGTGAGCGCTGAA |
| TGF-β2 | TCGACATGGATCAGTTTATGCG | CCCTGGTACTGTTGTAGATGGA |
| IL-1β | GAAATGCCACCTTTTGACAGTG | TGGATGCTCTCATCAGGACAG |
| TNF-α | CGTCAGCCGATTTGCTATCT | CGGACTCCGCAAAGTCTAAG |
| CX3CR1 | CTGTTATTTGGGCGACATTG | AACAGATTTCCCACCAGACC |
| CCR2 | CCTGCAAAGACCAGAAGAGG | GTGAGCAGGAAGAGCAGGTC |
| CXCL1 | TCCAGAGCTTGAAGGTGTTGCC | AACCAAGGGAGCTTCAGGGTCA |
| CX3CL-1 | CAGTGGCTTTGCTCATCCGCTA | AGCCTGGTGATCCAGATGCTTC |
