## Supplemental Table 2: List of Primary Antibodies. for "PAD2 knockout reduces myelin protein aggregates, modulates neuroinflammation and protects motor neurons, axons and neuromuscular junction in a SOD1-ALS mouse model"

**Table S2: List of Primary Antibodies.**

| Antibody | Company | Catalog No. | Dilution | RRID |
|---|---|---|---|---|
| PADI2 | Proteintech | 12110-1-AP | 1:1000 (WB)<br>1:50 (IF) | AB_2159475 |
| Anti-Modified citrulline, clone C4 (human recombinant) | Millipore | MABS487 | 1:1000 (WB)<br>1:100 (IF)<br>1:1000 (FTB) | |
| PADI3 (F-6) | Santacruz | sc-393622 | 1:100 (WB) | |
| GFAP | Cell Signaling | 3670S | 1:400 (IF) | AB_561049 |
| AIF-1/Iba1 | Novus Biologicals | NB100-1028 | 1:100 (IF) | AB_521594 |
| Anti-Iba1 | FUJIFILM Wako Pure Chemical Corporation | 019-19741 | 1:100 (IF) | AB_839504 |
| Anti-hOlig2 | R and D Systems | AF2418 | 1:100 (IF) | AB_2157554 |
| Anti-NFL | Zuoshang | | 1:300 (IF) | |
| NG2 | Millipore | AB5320 | 1:100 (IF) | AB_91789 |
| Myelin PLP | Novus Biologicals | NB100-1608 | 1:100 (IF)<br>1:1000 (WB)<br>1:1000 (FTB) | AB_2062362 |
| Anti-MBP, a.a. 82-87 | Millipore | MAB386 | 1:200 (IF)<br>1:1000 (WB)<br>1:1000 (FTB) | AB_94975 |
| MAG (A-11) | Santacruz | sc-166849 | 1:100 (IF)<br>1:1000 (WB)<br>1:1000 (FTB) | AB_2250078 |
| MOG (D-2) | Santacruz | sc-376138 | 1:100 (IF) | AB_10989782 |
| ChAT | Millipore | AB144P | 1:400 (IHC) | |
| CNPase | Cell Signaling | 2986S | 1:1000 (WB) | AB_2082474 |
| Anti-SOD1 | Non-commercial | BioDesign, Saco, ME, USA | 1:1000 (WB) | |
| CD68 | Biorad | MCA1957 | 1:200 (IF) | AB_322219 |
| ɣ-tubulin | Thermo Fisher Scientific | MA1-850 | 1:1000 (WB) | AB_2211249 |
| Synaptophysin | Thermo Fisher Scientific | MA5-16402 | 1:100 (IF) | AB_2537921 |
| Anti-Complement C3 | Novus Biologicals | NB200-540 | 1:100 (IF) | AB_10003444 |
| Alexa-488nm α-Bungarotoxin | Thermo Fisher Scientific | B13422 | 1:200 (IF) | |
| Ubiquitin | Dako | Z0458 | 1:1000 (FTB) | AB_2315524 |

FTB = Filter trap blot
