## Supplemental Table 3: List of Secondary Antibodies. for "PAD2 knockout reduces myelin protein aggregates, modulates neuroinflammation and protects motor neurons, axons and neuromuscular junction in a SOD1-ALS mouse model"

**Table S3: List of Secondary Antibodies.**

| Antibody | Catalog No. | Company | Dilution | RRID |
|---|---|---|---|---|
| Goat anti-Rabbit IgG (H+L), (HRP) | 65-6120 | Invitrogen | 1:5000 (WB) | AB_2533967 |
| Goat anti-Mouse IgG (H+L), (HRP) | 62-6520 | Thermo Fisher Scientific | 1:5000 (WB) | AB_2533947 |
| Rabbit anti-Sheep IgG, H & L Chain Specific HRP Conjugate | 402100 | Millipore | 1:5000 (WB) | AB_437820 |
| Goat Anti-Human IgG, HRP Conjugate | CS216591 | Millipore | 1:2000 (WB) | |
| Goat anti-Rat IgG (H+L), (HRP) | 31471 | Thermo Fisher Scientific | 1:5000 (WB) | AB_10965062 |
| Donkey anti-Chicken Igy (H+L) HRP Conjugate | SA1-300 | Thermo Fisher Scientific | 1:5000 (WB) | AB_325995 |
| Donkey anti-Goat IgG (H+L), (HRP) | A15999 | Invitrogen | 1:5000 (WB) | AB_2534673 |
| Alexa Fluor® 488-conjugated AffiniPure Goat Anti-Human IgG (H+L) | 109-545-003 | Jackson Immuno Research Laboratories | 1:250 (IF) | AB_2337831 |
| AffiniPure Goat Anti-Human IgG (H+L). Dylight™ 549 | 109-505-088 | Jackson Immuno Research Laboratories | 1:250 (IF) | AB_2337539 |
| Alexa Fluor™ 488 donkey anti-mouse IgG (H+L) | A21202 | Thermo Fisher Scientific | 1:250 (IF) | AB_141607 |
| Alexa Fluor® 568 donkey anti-mouse IgG (H+L) | A10037 | Thermo Fisher Scientific | 1:250 (IF) | AB_2534013 |
| Alexa Fluor™ 568 donkey anti-rabbit IgG (H+L) | A10042 | Thermo Fisher Scientific | 1:250 (IF) | AB_2534017 |
| Alexa Fluor® 568 goat anti-mouse IgG (H+L) | A11031 | Thermo Fisher Scientific | 1:250 (IF) | AB_144696 |
| Alexa Fluor™ 488 donkey anti-rabbit IgG (H+L) | A21206 | Thermo Fisher Scientific | 1:250 (IF) | AB_2535792 |
| Alexa Fluor® 488 goat anti-rabbit IgG (H+L) | A11034 | Thermo Fisher Scientific | 1:250 (IF) | AB_2576217 |
| Alexa Fluor® 594 goat anti-rabbit IgG (H+L) | A11037 | Thermo Fisher Scientific | 1:250 (IF) | AB_2534095 |
| Alexa Fluor® 488 goat anti-mouse IgG (H+L) | A11029 | Thermo Fisher Scientific | 1:250 (IF) | AB_2534088 |
| Alexa Fluor® 568 donkey anti-goat IgG (H+L) | A11057 | Thermo Fisher Scientific | 1:250 (IF) | AB_2534104 |
