## Supplemental Figure 1: Genetic knockout of PAD2 eliminates PAD2 expression. for "PAD2 knockout reduces myelin protein aggregates, modulates neuroinflammation and protects motor neurons, axons and neuromuscular junction in a SOD1-ALS mouse model"

Fig S1

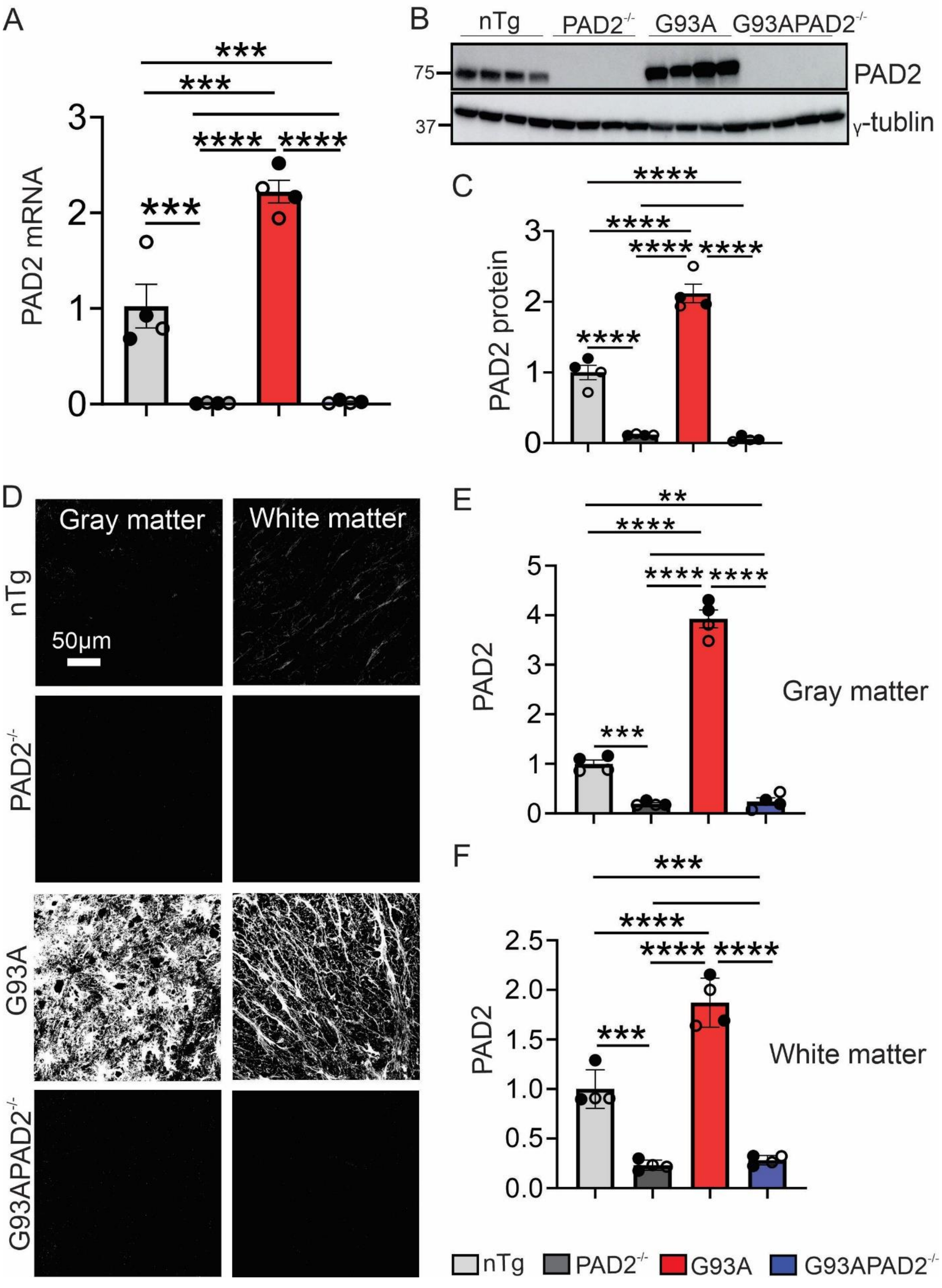
A
PAD2 mRNA
***
****
B
nTg
PAD2-/-
G93A
G93APAD2-/-
75
37
PAD2
γ-tublin
C
PAD2 protein
D
Gray matter
White matter
nTg
PAD2-/-
G93A
G93APAD2-/-
50μm
E
PAD2
Gray matter
F
PAD2
White matter
nTg
PAD2-/-
G93A
G93APAD2-/-

**Figure S1. Genetic knockout of PAD2 eliminates PAD2 expression.** (A) A qPCR for quantification of PAD2 mRNA. (B) A PAD2 protein blot. (C) Quantification of PAD2 protein levels in B. (D) Immunofluorescence staining for PAD2 in ventral horn and white matter. (E, F) Quantification of fluorescence intensity of PAD2 in ventral horn and white matter, respectively. The experiments were performed on lumber spinal cords at disease end-stage of $SOD1^{G93A}$ mice. n = 4 per group. Each circle represents one mouse. Filled circles represent males and unfilled circles represent females. Statistics: one-way ANOVA with Bonferroni post-hoc test. *$p<0.05$. **$p<0.01$. ***$p<0.001$. ****$p<0.0001$.
