## Supplemental Figure 2: PAD2 knockout does not alter other expressions of other PADs. for "PAD2 knockout reduces myelin protein aggregates, modulates neuroinflammation and protects motor neurons, axons and neuromuscular junction in a SOD1-ALS mouse model"

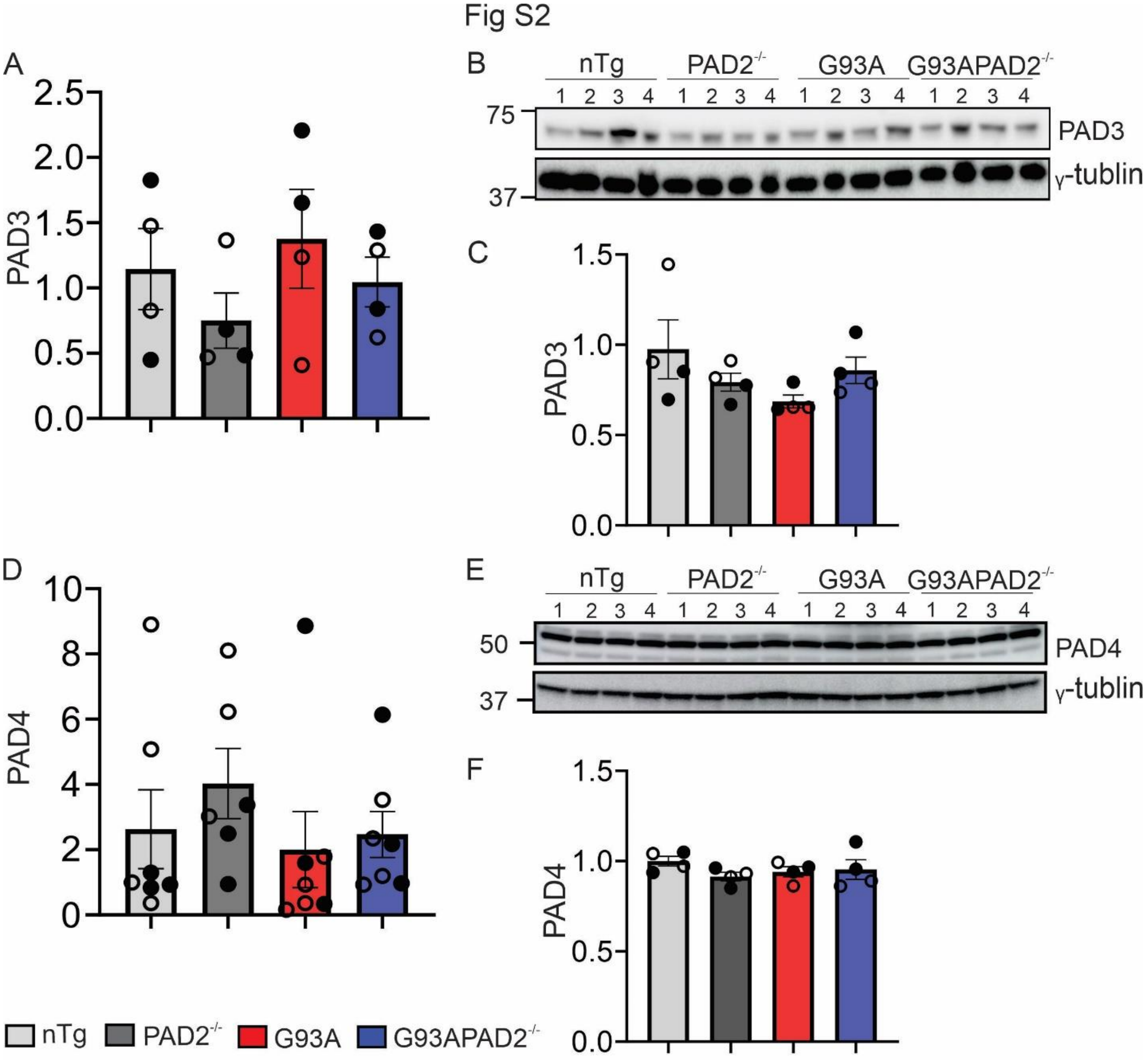


**Figure S2. PAD2 knockout does not alter other expressions of other PADs.** (A) Levels of PAD3 mRNA in mouse lumber spinal cords by qPCR. (B) Western blot for PAD3 in lumber spinal cords. (C) Quantifications of PAD3 signal density in B. (D) Levels of PAD4 mRNA in mouse lumber spinal cords by qPCR. (E) Western blot for PAD4 in lumber spinal cords. (F) Quantifications of PAD4 signal density in E. n = 4-8 per group. Symbols and statistics are as described in Figure S1.
