## Supplemental Figure 3: Current commercial PAD4 antibodies are not specific for PAD4 in mouse spinal cord lysate. PAD4 antibodies, including. for "PAD2 knockout reduces myelin protein aggregates, modulates neuroinflammation and protects motor neurons, axons and neuromuscular junction in a SOD1-ALS mouse model"

Fig. S3

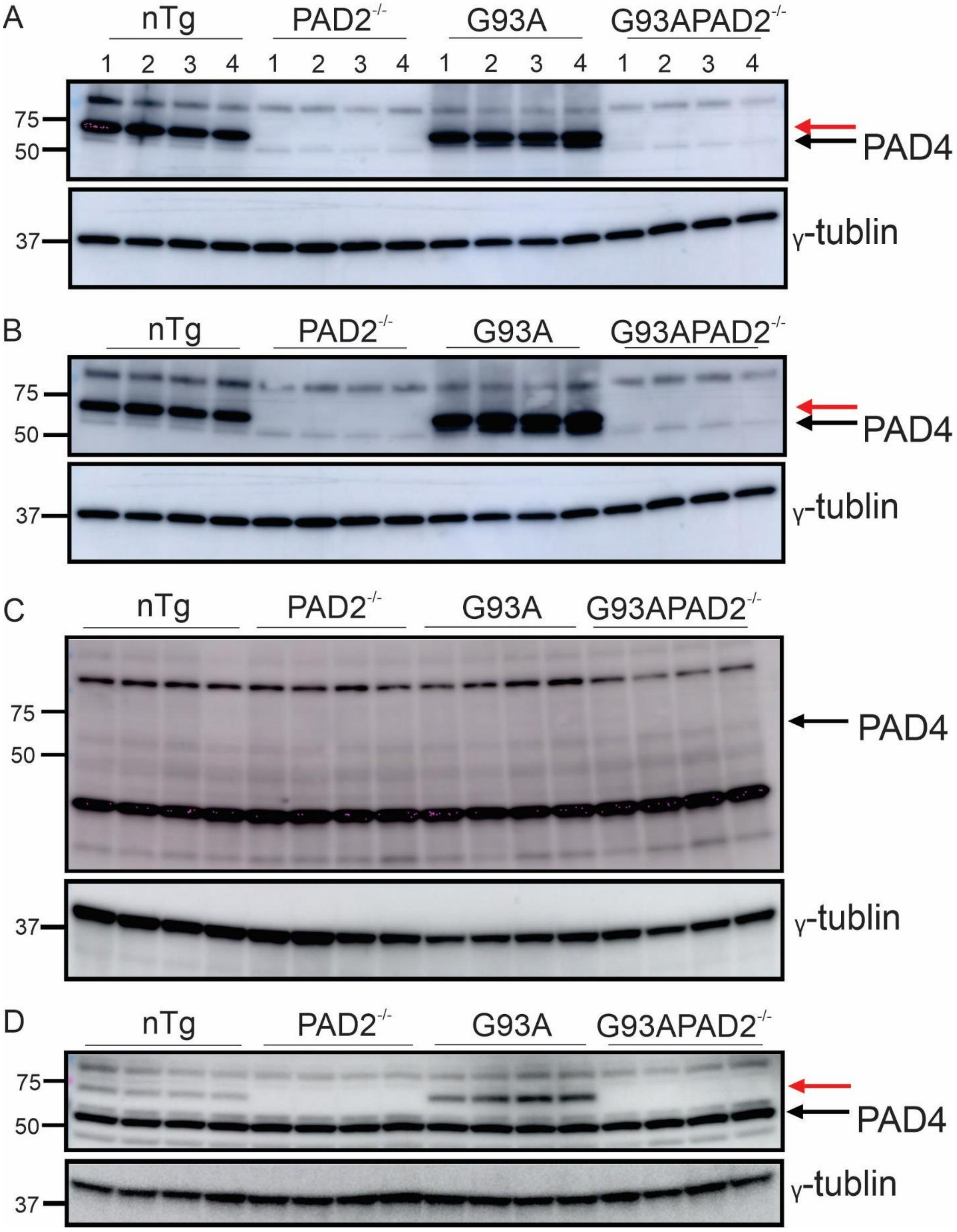

A
nTg
PAD2-/-
G93A
G93APAD2-/-
1 2 3 4 1 2 3 4 1 2 3 4 1 2 3 4
75
50
PAD4
37
γ-tublin
B
nTg
PAD2-/-
G93A
G93APAD2-/-
75
50
PAD4
37
γ-tublin
C
nTg
PAD2-/-
G93A
G93APAD2-/-
75
50
PAD4
37
γ-tublin
D
nTg
PAD2-/-
G93A
G93APAD2-/-
75
50
PAD4
37
γ-tublin

**Figure S3. Current commercial PAD4 antibodies are not specific for PAD4 in mouse spinal cord lysate. PAD4 antibodies, including.** (A) Abcam AB96758 polyclonal antibody, (B) Abcam AB50247 polyclonal antibody, (C) Abcam AB128086 monoclonal antibody, and (D) PAD4 Proteintech 17373-1-AP antibody, were tested by Western blots with mouse spinal cord lysates. In addition to PAD4, three of the four reacted with PAD2 and all reacted with other non-PAD4 proteins. Red arrows point to strong PAD2 signal. Black arrows point to PAD4 signal.
