## Supplemental Figure 4: PAD2 does not contribute to SOD1 and ubiquitin aggregation in SOD1G93A mice. for "PAD2 knockout reduces myelin protein aggregates, modulates neuroinflammation and protects motor neurons, axons and neuromuscular junction in a SOD1-ALS mouse model"

Fig S4

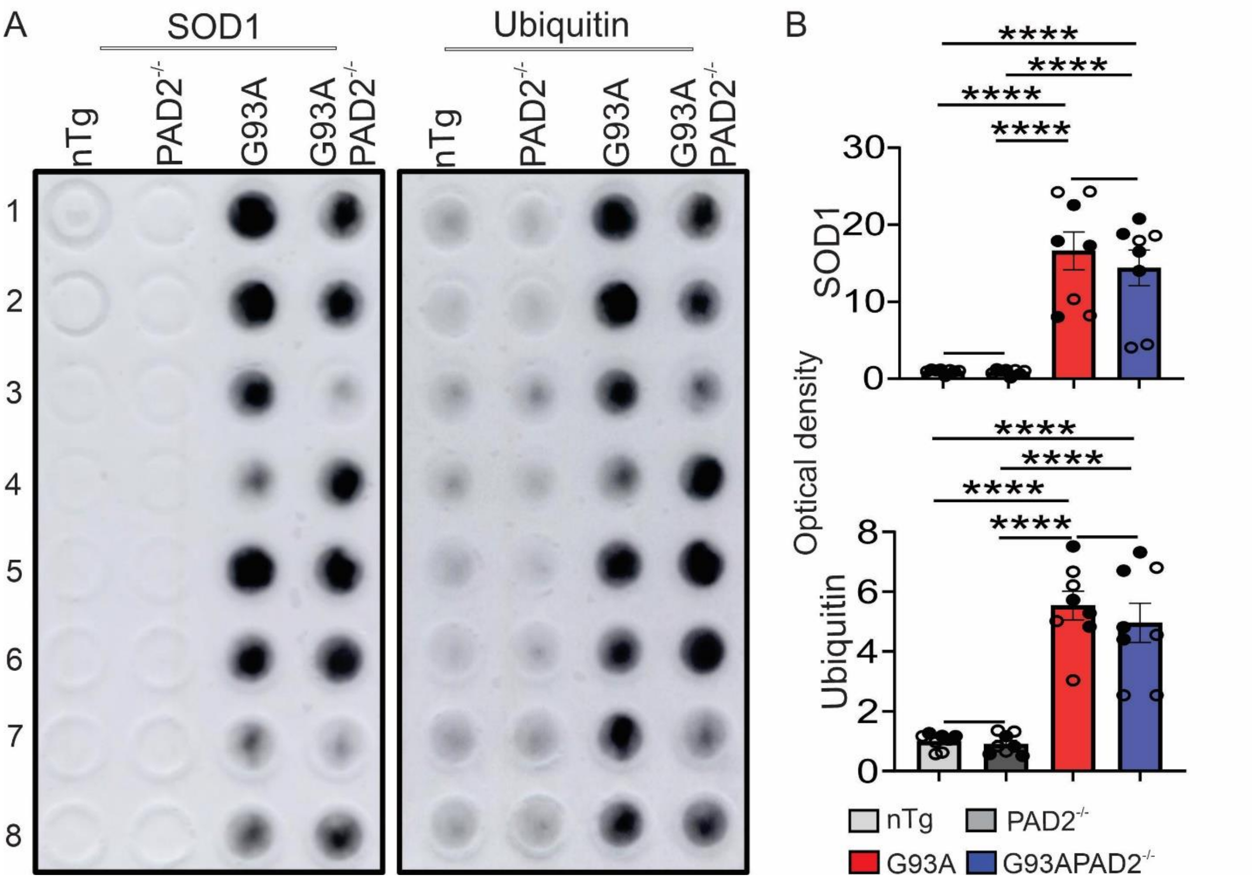


**Figure S4. PAD2 does not contribute to SOD1 and ubiquitin aggregation in SOD1$^{G93A}$ mice.** (A) Filter trap assay for protein aggregates in mouse spinal cord. (B) Quantification of optical density of SOD1 and ubiquitin in (A). n = 8 in all groups. Symbols and statistics are as described in Figure S1.
