## Supplemental Figure 5: PAD2 knockout does not alter myelin gene expression in SOD1G93A mice. for "PAD2 knockout reduces myelin protein aggregates, modulates neuroinflammation and protects motor neurons, axons and neuromuscular junction in a SOD1-ALS mouse model"

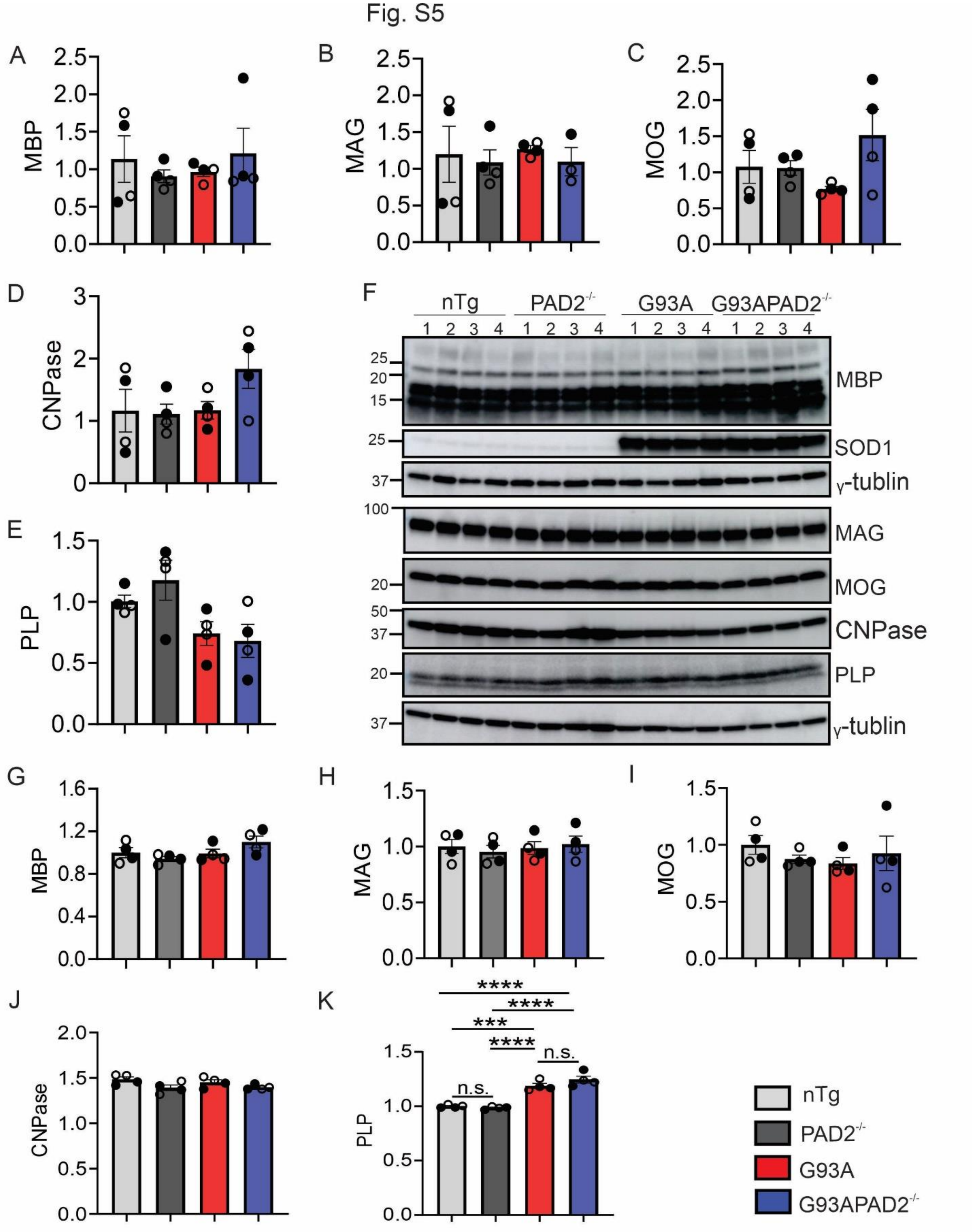


**Figure S5. PAD2 knockout does not alter myelin gene expression in SOD1$^{G93A}$ mice.** (A-E) Levels of MBP, MAG, MOG, CNPase, and PLP mRNA transcripts from lumber spinal cords. (F)

Western blots for myelin protein levels lumber spinal cords. (G-K) Quantifications of band signal density of MBP, MAG, MOG, CNPase, and PLP in (F). n = 4 in all groups. Symbols and statistics are described in Figure S1.
