## Supplemental Figure 6: PAD2 knockout does not impact gliosis in the spinal cord of SOD1G93A mice. for "PAD2 knockout reduces myelin protein aggregates, modulates neuroinflammation and protects motor neurons, axons and neuromuscular junction in a SOD1-ALS mouse model"

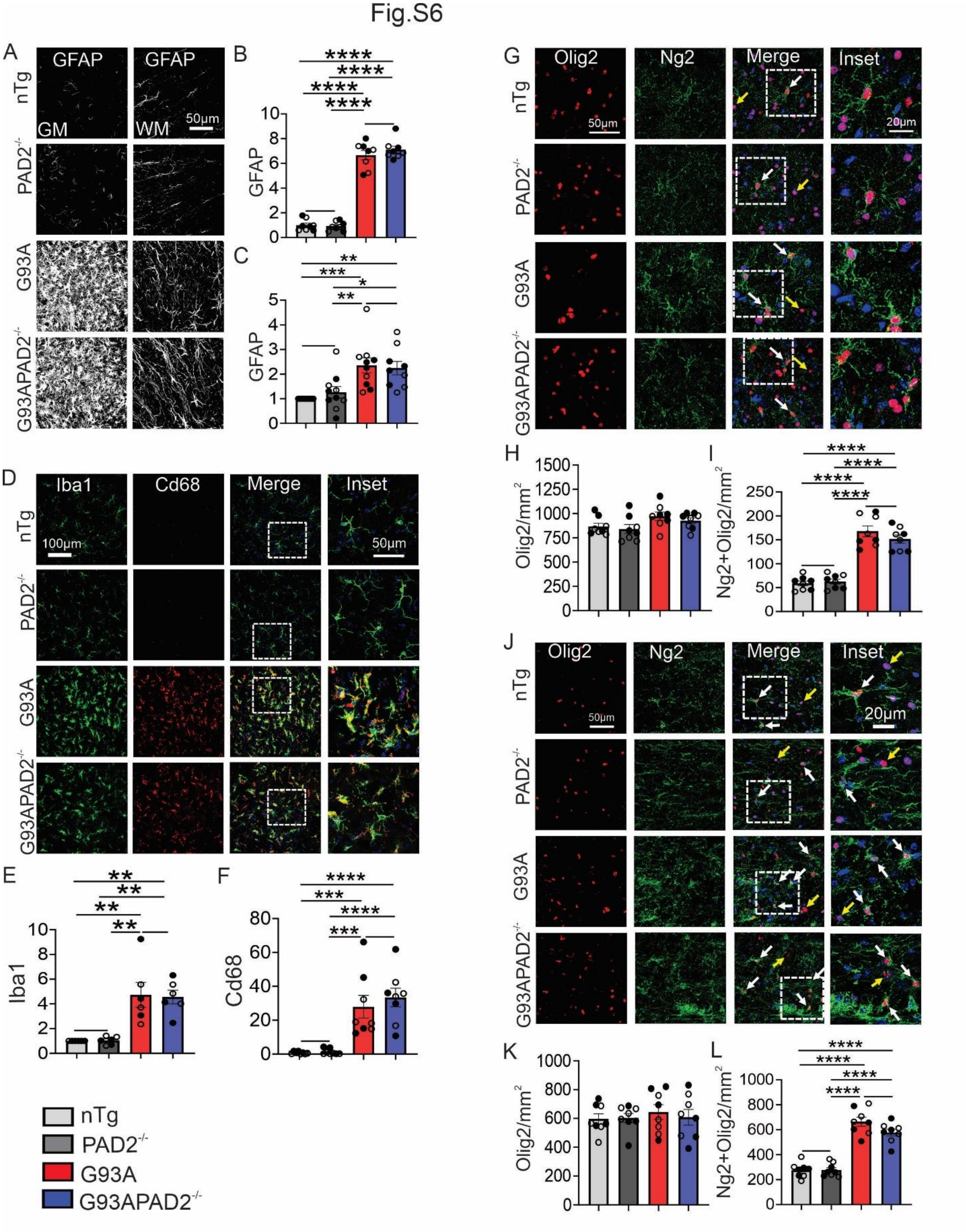


**Figure S6. PAD2 knockout does not impact gliosis in the spinal cord of SOD1$^{G93A}$ mice.** (A) Immunofluorescence staining for GFAP in the ventral (GM) and white matter (WM). (B, C)

Quantification of fluorescent intensity of GFAP in the gray and white matter respectively, as shown in A. (D) Double immunofluorescence staining for Iba1 and CD68 in ventral gray matter. (E, F) Quantification of fluorescent intensity of Iba1 and CD68 in ventral gray matter, respectively, from images as shown in D. (G) Double immunofluorescence staining for Olig2 and Ng2 in ventral gray matter. (H, I) Quantification of Olig2 positive cells, and Ng2 positive-Olig2 positive cell numbers in ventral gray matter, respectively in G. (J) Double immunofluorescence staining for Olig2 and Ng2 in spinal cord white matter. White arrows point to Ng2 positive Olig2 positive cells. Yellow arrows point to Ng2 negative Olig2 positive cells. (K, L) Quantification of Olig2 positive cells and of Ng2 positive-Olig2 positive cell numbers, respectively, from images as shown in J. n = 6-8 in each group. Symbols and statistics are described in Fig. S1.
