## Supplemental Figure 7: PAD2 knockout does not impact the expression of many inflammatory markers in the spinal cord of SOD1G93A mice. for "PAD2 knockout reduces myelin protein aggregates, modulates neuroinflammation and protects motor neurons, axons and neuromuscular junction in a SOD1-ALS mouse model"

Fig.S7

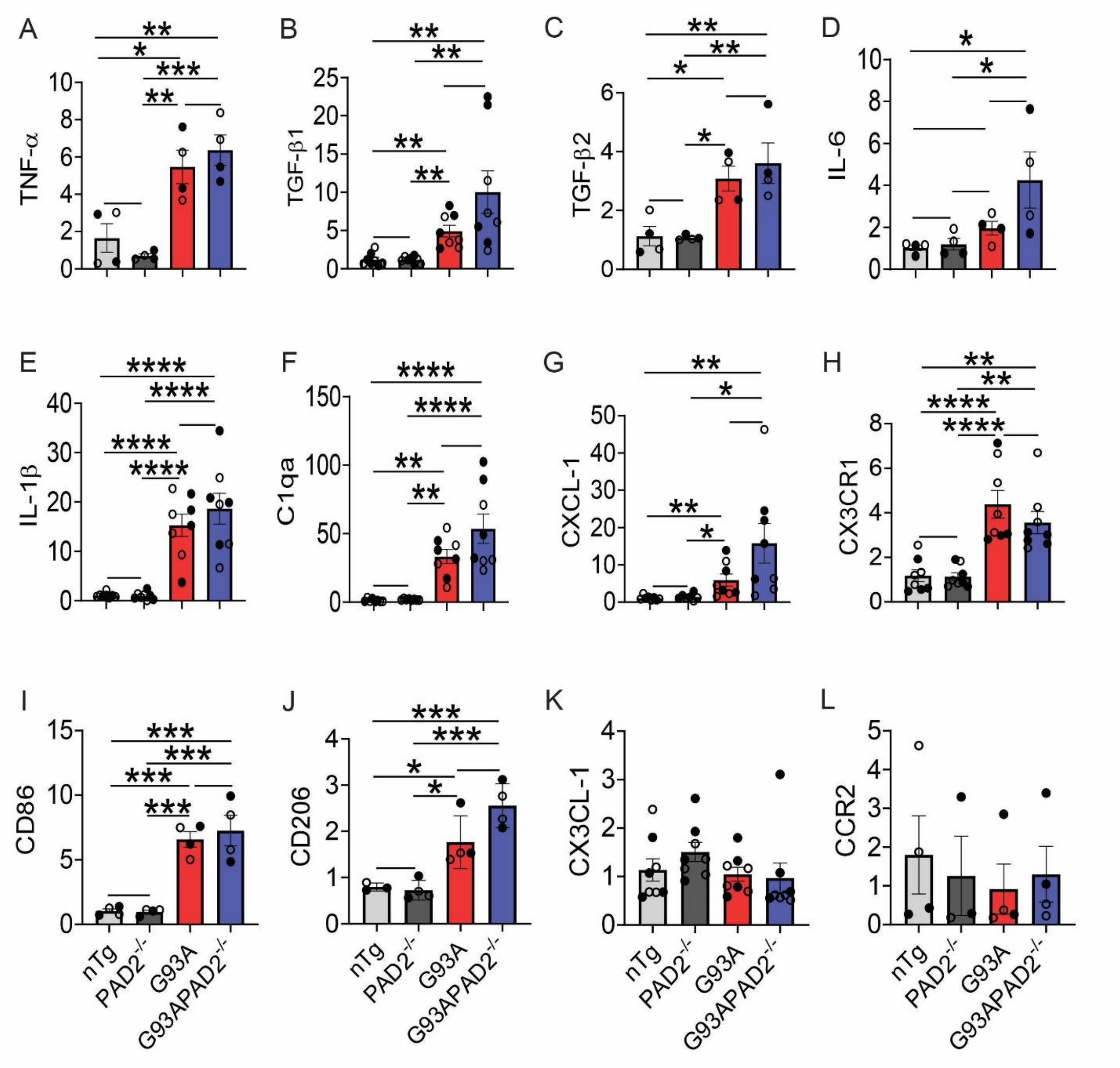


**Figure S7. PAD2 knockout does not impact the expression of many inflammatory markers in the spinal cord of SOD1$^{G93A}$ mice.** (A-L) A qPCR for quantification of inflammatory markers including TNF-α, TGF-β1, TGF-β2, IL-6, IL-1β, C1qa, CXCL-1, CX3CR1, CD86, CD206, CX3CL-1, CCR2 mRNAs, respectively.
