## Supplemental Figure 8: PAD2 knockout reduces C3 in astrocytes in the spina cord gray matter in SOD1G93A mice. for "PAD2 knockout reduces myelin protein aggregates, modulates neuroinflammation and protects motor neurons, axons and neuromuscular junction in a SOD1-ALS mouse model"

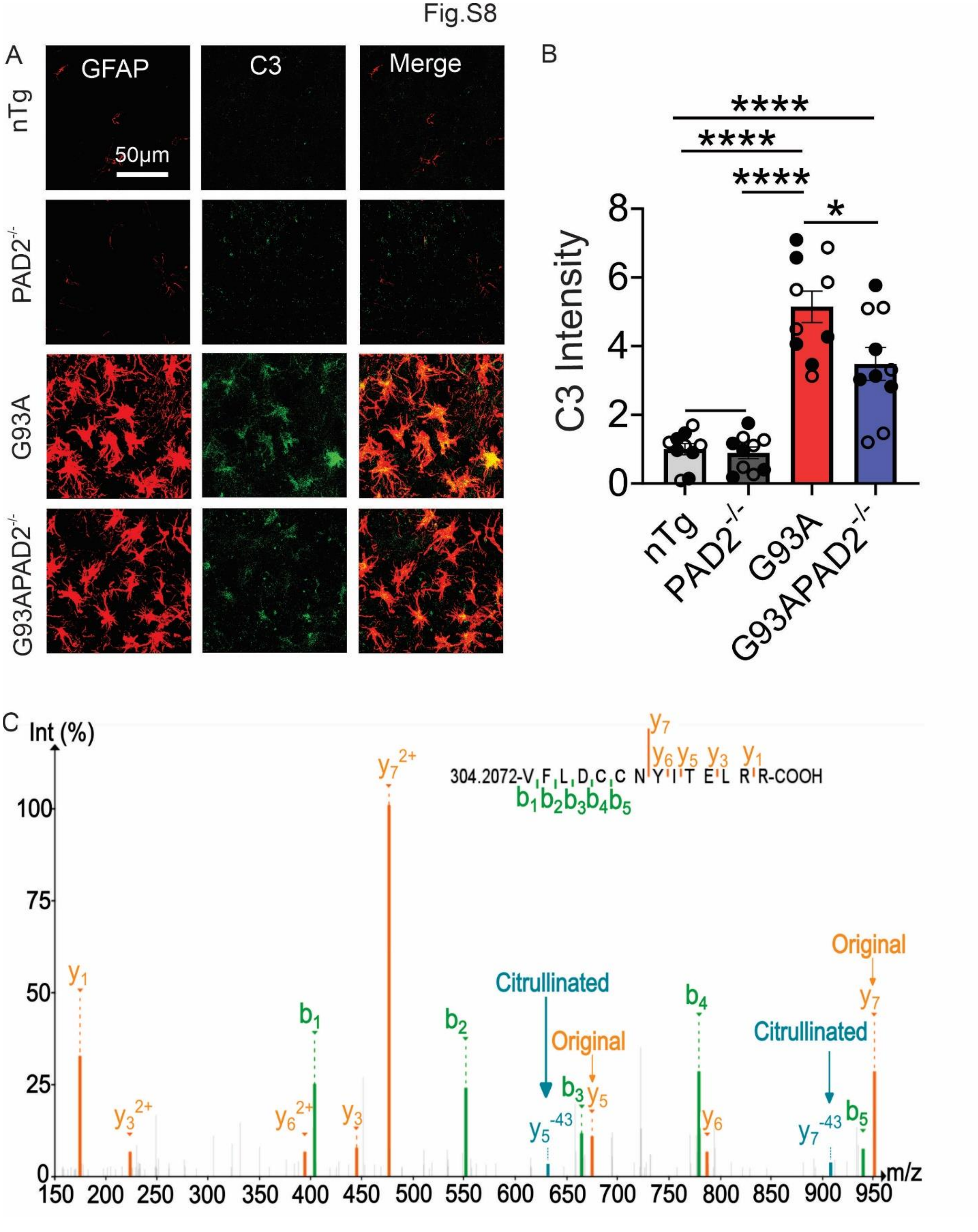


**Figure S8. PAD2 knockout reduces C3 in astrocytes in the spina cord gray matter in SOD1$^{G93A}$ mice.** (A) Double immunofluorescence staining for GFAP and C3. (B) Quantification of C3 intensity from images as shown in A. n = 10 in each group. Symbols and statistics are described in Fig. S1. (C) Fragmentation spectra from the peptide containing R735cit in C3 identified by ionFinder. The diagnostic spectra for citrullination with the neutral loss (-43Da) are indicated. The panel was generated by Proteomics Data Viewer (PDV) software.
