## Supplemental Table 4: The onset and lifespan medians, averages and statistics for "PAD2 knockout reduces myelin protein aggregates, modulates neuroinflammation and protects motor neurons, axons and neuromuscular junction in a SOD1-ALS mouse model"

**Table S4: The onset and lifespan medians, averages and statistics**

| | | Male | | | | | Female | | | |
|---|---|---|---|---|---|---|---|---|---|---|
| | | Average | Std | Median | P value | | Average | Std | Median | P value |
| **Lifespan (days)** | **SOD1$^{G93A}$** | 147.3 | 11.2 | 144 | *0.0261 | | 158.1 | 14.7 | 149 | 0.8597 |
| | **SOD1$^{G93A}$/ PAD2KO** | 154.6 | 18.4 | 158 | | | 158.1 | 23.3 | 155 | |
| **Onset (weeks)** | **SOD1$^{G93A}$** | 15.25 | 2.527 | 15 | 0.1924 | | 16.69 | 2.463 | 17 | 0.2394 |
| | **SOD1$^{G93A}$/ PAD2KO** | 14.07 | 1.94 | 14 | | | 15.07 | 2.67 | 14 | |
| **Disease duration (weeks)** | **SOD1$^{G93A}$** | 6.875 | 2.715 | 6 | 0.2466 | | 6.385 | 3.015 | 6 | 0.406 |
| | **SOD1$^{G93A}$/ PAD2KO** | 8.133 | 3.08 | 7 | | | 7.786 | 2.49 | 8 | |
| | **SOD1$^{G93A}$** | n=12 | | | | | n=13 | | | |
| | **SOD1$^{G93A}$/ PAD2KO** | n=15 | | | | | n=14 | | | |
