## Supplemental Table 5: Onset and survival data from individual mice for "PAD2 knockout reduces myelin protein aggregates, modulates neuroinflammation and protects motor neurons, axons and neuromuscular junction in a SOD1-ALS mouse model"

**Table S5: Onset and survival data from individual mice**

| | SOD1$^{G93A}$ | | | | SOD1$^{G93A}$/PAD2KO | | | |
|---|---|---|---|---|---|---|---|---|
| | Male | | Female | | Male | | Female | |
| | Onset (weeks) | Survival (days) | Onset (weeks) | Survival (days) | Onset (weeks) | Survival (days) | Onset (weeks) | Survival (days) |
| 1 | 12 | 140 | 18 | 137 | 14 | 137 | 14 | 125 |
| 2 | 15 | 141 | 11 | 147 | 13 | 139 | 14 | 131 |
| 3 | 16 | 143 | 18 | 149 | 13 | 139 | 12 | 138 |
| 4 | 16 | 146 | 18 | 150 | 15 | 144 | 14 | 139 |
| 5 | 14 | 152 | 17 | 153 | 13 | 144 | 12 | 148 |
| 6 | 15 | 153 | 17 | 153 | 16 | 146 | 17 | 148 |
| 7 | 17 | 156 | 17 | 167 | 12 | 148 | 14 | 155 |
| 8 | 13 | 164 | 12 | 167 | 14 | 149 | 13 | 163 |
| 9 | 19 | 165 | 19 | 172 | 11 | 151 | 15 | 166 |
| 10 | 14 | 167 | 17 | 172 | 16 | 154 | 16 | 171 |
| 11 | 20 | 168 | 16 | 175 | 17 | 158 | 15 | 180 |
| 12 | 12 | 170 | 19 | 178 | 17 | 167 | 20 | 181 |
| 13 | | | 18 | 186 | 14 | 176 | 14 | 181 |
| 14 | | | | | 11 | 186 | 21 | 208 |
| 15 | | | | | 15 | 199 | | |
